# Integrated cell atlas and tumoroids chart pancreatic cancer therapeutic targets

**DOI:** 10.1101/2025.09.30.679518

**Authors:** Quan Xu, Ryo Okuda, Bruno Gjeta, Christoph Harmel, Marina Signer, Matilde Lucioli, Malgorzata Santel, Makiko Seimiya, Cinzia Esposito, Karolina Guja-Jarosz, Ashley Maynard, Soichiro Morinaga, Yohei Miyagi, Tomoyuki Yamaguchi, Yasuharu Ueno, Salvatore Piscuoglio, Daniel J. Müller, Hideki Taniguichi, Barbara Treutlein, J. Gray Camp

**Author notes:** Correspondence: HT; BT; JGC. These authors contributed equally (Quan Xu, Ryo Okuda, Bruno Gjeta).

## Abstract

Pancreatic ductal adenocarcinoma (PDAC) is characterized by dense, fibroblast-rich stroma that actively shapes the tumor microenvironment. Most PDAC cases arise from conserved genetic transformations initiated by oncogenic KRAS mutations, developing into metastatic disease with high mortality rates. To chart universal PDAC cell states and identify therapeutic inroads, we integrated published single-cell transcriptomes from 200 patient samples, and used the atlas to define prevalent cancer cell and cancer-associated fibroblast (CAF) states, gene expression programs, and ligand-receptor interactions. We established modular tumoroids incorporating patient-derived cancer cells and CAFs that recapitulate aspects of ductal architecture and desmoplastic stroma. Single-cell and spatial transcriptomic profiling confirmed preservation of key cellular states and signaling networks in vitro. We identified Syndecan-1 (SDC1) as a CAF-responsive cancer cell receptor correlating with poor patient survival. Functional SDC1 blockade disrupted cancer growth in tumoroids, highlighting therapeutic relevance. This study provides a framework for dissecting cancer-stroma dynamics and identifying actionable targets using patient-derived tumoroid models.

## Introduction

Pancreatic ductal adenocarcinoma (PDAC) is one of the most aggressive and intractable forms of cancer (Siegel et al., 2025). PDAC tumors at diagnosis are characterized by stark intra-tumoral heterogeneity with a dense stroma component which can constitute over 70% of the tumor mass. Inhibition of oncogenic KRAS is an early causal genetic lesion in most (90%) PDAC tumors, and subsequent evolutionary events are shaped by the particular tumor microenvironment (TME) within the pancreas (Halbrook et al., 2023; Dreyer et al., 2025). The pancreas is divided into four main anatomical regions (head, neck, and body tail), and PDAC can arise from ductal epithelial cells located within each one of these regions. The pancreatic ductal epithelial origin and the general microenvironment of the pancreas likely place similar evolutionary pressures on PDAC development across patients, however it is unclear if there are universal states that occur in the majority of PDAC indications (Halbrook et al., 2023; Burdziak et al., 2023).

Cancer associated fibroblasts (CAFs) are central PDAC stroma components that coordinate the TME by building an extracellular matrix and secreting cytokines and growth factors that regulate cancer growth (Sahai et al., 2020). Microenvironmental pressures including hypoxia and nutrient scarcity within the primary tumor can lead individual cancer cells to acquire specific metabolic and other cell state signatures that support cancer cell adaptation to current conditions, and also provide the ability for future colonization into other organ niches (Massey et al., 2024; Mo et al., 2024; Agrawal et al., 2025). However, a thorough understanding of how CAF-cancer cell interactions give rise to diverse tumor cell states remains incomplete, due in part to the spatial and temporal complexity of these interactions in vivo.

To dissect the PDAC tumor ecosystem, there is a need for integrative resources that unify cellular states and intercellular signaling across patients. Single-cell transcriptomics provides a powerful approach for resolving cellular heterogeneity, reconstructing developmental hierarchies, and deciphering cell-cell communication in both health and disease (Heumos et al., 2023; Skinnider et al., 2025; He et al., 2024; Xu et al., 2025). Integration of single-cell PDAC data could enable the identification of commonly observed cancer and stromal cell states and their interaction networks across patients, serving as a reference for mechanistic studies. In parallel, patient-derived in vitro models such as cancer cystic organoids (CCOs) offer a tractable platform to study tumor biology in a controlled setting (Verstegen et al., 2025; Mizutani et al., 2024; Dayton et al., 2023; Boj et al., 2015). When combined with co-cultured stromal components, such as CAFs, these models can recapitulate relevant aspects of the TME (Öhlund et al., 2017; Seino et al., 2018; Kim et al., 2020; Barbazan et al., 2023). Here, we set out to build a reference PDAC cell atlas and use it to benchmark a modular, stroma-rich tumoroid co-culture system. Through this combined strategy, we uncover principles of PDAC cell-state diversity, intercellular communication, and potential therapeutic targets within the tumor microenvironment.

## Results

### An integrated PDAC single-cell transcriptome atlas

To characterize the gene expression patterns of human PDAC cells, we compiled single-cell RNA sequencing (scRNA-seq) data from 12 published datasets, encompassing 673,000 cells from 200 samples (Fig. 1a, Extended Data Table 1). These datasets were generated using multiple sequencing protocols, primarily commercialized droplet-based methods (Fig. 1). For some datasets, paired cancer and healthy tissue samples were available and annotated. We performed high-resolution clustering within each dataset and assigned cell-type annotations based on known marker gene expression and differential expression analysis between clusters. To enable consistent, label-aware integration, we implemented a two-level hierarchical annotation system, defining broad cell classes (level 1) and refined cell types (level 2) (Fig. 1). To address batch effects and achieve robust atlas integration, we evaluated 12 different data integration methods using single-cell integration benchmarking (Polański et al., 2020; Xu et al., 2021; Lopez et al., 2018; Korsunsky et al., 2019; He et al., 2020; Büttner et al., 2019; De Donno et al., 2023; Hao et al., 2024) (Extended Data Fig. 1). Based on this assessment, we selected scPoli (Luecken et al., 2022; De Donno et al., 2023), which has been shown to effectively integrate heterogeneous single-cell datasets (He et al., 2024; Xu et al., 2025), to generate an integrated embedding of all PDAC cells, providing a cohesive representation of the diverse datasets (Fig. 1b). The resulting PDAC cell atlas encompasses seven major tissue-level cell classes and 28 refined level 2 cell types (Fig. 1b, Extended Data Fig. 2a,b). Comparison of integration cell-type annotations with those reported in the original studies showed high concordance for most cell labels (Extended Data Fig. 2c,d) (Chijimatsu et al., 2022; Storrs et al., 2023). Overall, the integration was largely unaffected by patient metadata such as gender, age, or study origin (Fig. 1c). However, cell type composition varied across samples, particularly with respect to sample type (tumor versus adjacent normal) and tumor stage (Fig. 1d). To investigate cell-cell communication among PDAC cell types, we performed ligand-receptor pairing analysis using the integrated atlas. This analysis revealed a ligand-receptor interaction network spanning all major cell types. Notably, mesenchymal cells, Schwann cells, endothelial cells, and epithelial cells exhibited a higher number of interactions with other cell populations, suggesting active roles in intercellular signaling (Palikuqi et al., 2020; Thiel et al., 2025; Camp et al., 2017). In contrast, lymphoid, mast, endocrine, and erythrocyte cells showed relatively fewer interactions (Extended Data Fig. 2e,f).

**Figure 1.**
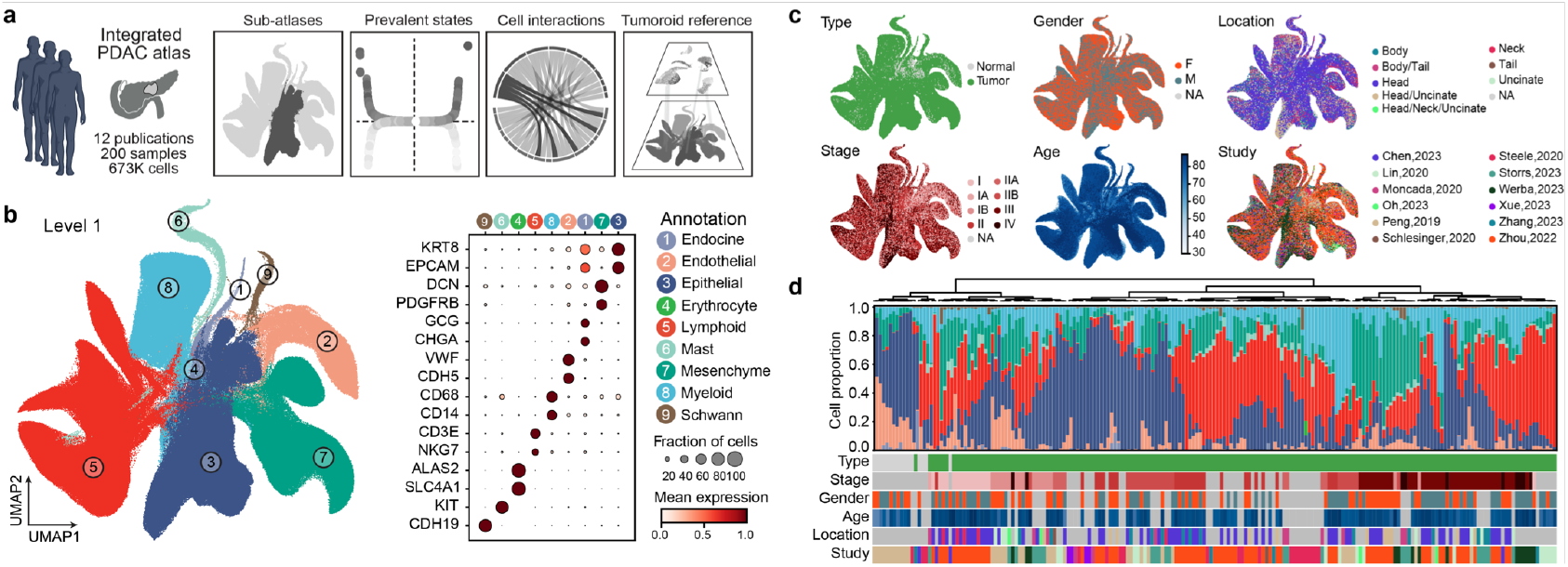
An integrated PDAC single-cell transcriptome atlas. (a) PDAC atlas integration combines 12 published datasets comprising 200 samples and 673,000 cells to identify the most prevalent gene expression states and cell-cell interactions across patients. (b) Left, UMAP embedding of the integrated atlas with Level 1 annotations, shows eight major cell types: lymphoid, endocrine, endothelial, mesenchymal, epithelial, myeloid, erythrocytes, and Schwann cells. Right, Dot plot shows marker expression for each major cell type. (c) UMAP embedding of all cells, colored by sample metadata. From left to right and top to bottom, the panels represent sample type, gender, anatomical location, tumor stage, age group, and study origin. (d) Stacked bar plot showing the proportion of cell types in each sample. Sidebars indicate sample annotations corresponding to the metadata shown in (c).

### Prevalent PDAC cancer-fibroblast interactions associate with patient survival

To characterize epithelial and mesenchymal cell heterogeneity in PDAC, we extracted and integrated epithelial and mesenchymal cells separately from the PDAC atlas (Fig. 2a-c, Extended Data Fig. 3a-c). The epithelial cells clustered into 12 distinct subtypes, including two normal cell types abundant in the non-cancerous tissue counterparts (Acinar (CTRB2) and Ductal(AQP1)), two pre-cancerous subtypes (Pre-cancer (BMX) and Precancer (MUC5B)), four classical-like cancer subtypes (Classical (CRABP2), Classical (CLDN18), Classical (SLC40A1), and Classical (CTTN)), and three basal-like cancer subtypes (Basal (KRT5), Basal (S100A4), and Basal (KRT7)) (Fig. 2b). For the mesenchymal compartment, we identified nine distinct subtypes, including one normal cell type (Stellate (MACM)), one pre-CAF sub-type (Pre-CAF (TAGLN2)), two complement-secreting CAFs (csCAF) (csCAF (CFH) and csCAF (C7)), two myofibroblast CAFs (myCAF) (myCAF (MMP11) and myCAF (HOPX)), one antigen-presenting CAF (apCAF) (apCAF (CD74)), and two inflammatory CAFs (iCAF) (iCAF (SOD2) and iCAF (EMP1)) (Fig. 2c).

**Figure 2.**
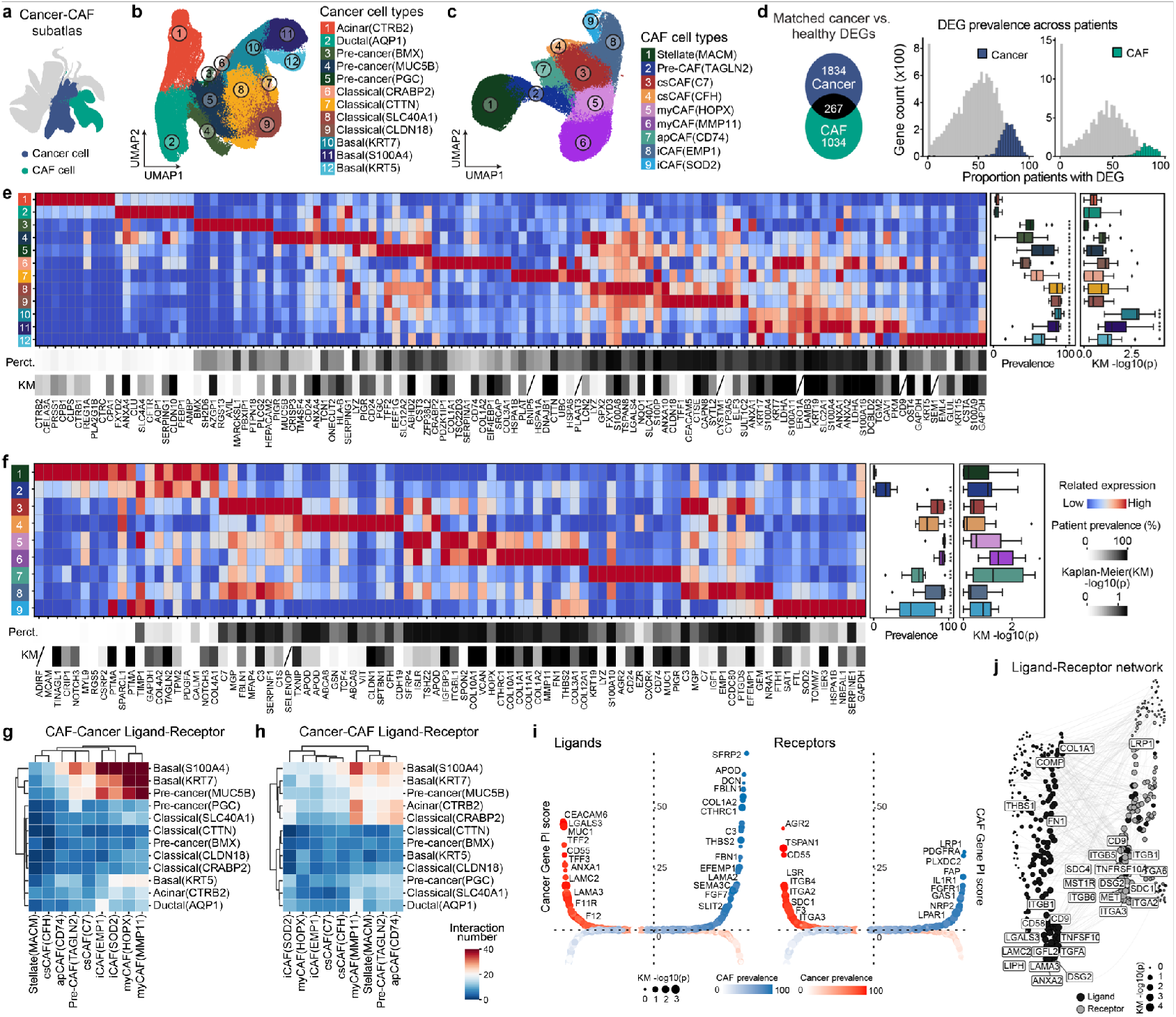
Highly prevalent cancer and CAF cell states interact and associate with survival. (a) UMAP embedding of the integrated atlas highlighting cancer and CAF (cancer-associated fibroblast) cells. (b) UMAP of epithelial cell subtypes identifying 12 distinct cancer cell populations, including acinar, ductal, pre-cancerous, classical, and basal-like states. (c) UMAP of mesenchymal cell subtypes revealing 10 distinct CAF populations, including stellate cells, pre-CAFs, cancer-stimulated CAFs (csCAFs), myofibroblastic CAFs (myCAFs), antigen-presenting CAFs (apCAFs), and inflammatory CAFs (iCAFs). (d) Venn diagram showing differentially expressed genes specific to cancer cells, CAFs, and those shared between both populations. Right: histograms showing the prevalence of each gene across patients, indicating the percentage of patients in which each gene is expressed in cancer cells (top) and CAFs (bottom). (e) Heatmap showing the top 10 marker genes for each cancer cell subtype. The top scale bar indicates the percentage of patients in which each gene is highly expressed; the bottom scale bar represents the −*log*_10_(*p*) value from Kaplan-Meier survival analysis. Genes not present in the TCGA database are marked with a black slash. Right: boxplots summarizing, from left to right, the patient prevalence and the −*log*10 (*p*) values of marker genes for each cancer cell subtype (^*^p < 0.05, ^**^p < 0.01, ^***^p < 0.001). (f) Same as (e), but for CAF cell subtypes. (g) Heatmap of inferred ligand-receptor interactions from cancer cells to CAFs in primary tissues. (h) Heatmap of inferred ligand-receptor interactions from CAFs to cancer cells in primary tissues. (i) Dot plots of all ligand (left) and receptor (right) genes. Genes are sorted left-to-right by cancer gene PI scores and right-to-left by CAF gene PI scores. Color indicates the percentage of samples with high gene expression, and dot size reflects the −*log*_10_(*p*) from survival analysis. (j) Ligand-receptor interaction network between cancer cells and CAFs. Dot size represents the −*log*10 (*p*) from survival analysis, with the top significant genes highlighted.

To further assess PDAC-associated features across patients, we merged all normal epithelial cells with cancer cells, and normal mesenchymal cells with CAFs, grouping them by sample. We then performed pairwise Wilcoxon tests for each gene to identify significantly differentially expressed genes. This analysis resulted in 1,834 cancer-specific genes, 1,043 CAF-specific genes, and 267 genes shared between both cancer and CAF populations that were consistently expressed across all patients (Fig. 2d, Extended Data Table 2). Pathway analysis revealed enrichment of EGF/EGFR, TGF, ErbB, and adhesion-related genes in the cancer-specific set; Wnt, focal adhesion, and PI3K pathway genes in CAF-specific genes; and metabolic and immune infiltration-related genes in the shared set (Extended Data Fig. 3d,e). Next, to characterize the expression patterns of the identified cancer and CAF cell types, we selected the top 10 feature genes for each cell type (Fig. 2e,f). The cancer cell and CAF feature genes exhibit a higher percentage of patient expression compared to the feature genes of normal epithelial and mesenchymal cells (Fig. 2e,f). To evaluate the relationship between these cell types and patient survival, we analyzed Kaplan-Meier survival curves using gene expression data from The Cancer Genome Atlas (TCGA) PDAC cohort, focusing on the expression of the identified feature genes. Notably, feature genes associated with three basal-like cancer cell subtypes showed a strong correlation with poor patient survival (Fig. 2e). Similarly, feature genes linked to myCAF (MMP11) also correlated with worse survival outcomes (Fig. 2e,f). These findings suggest that basal-like cancer cells and myCAFs represent divergent and potentially more aggressive cellular states in PDAC (Pei et al., 2025).

We analyzed ligands, receptors and other components of the microenvironment using the integrated atlas data. Basal-like cancer cells exhibited more extensive interactions with CAF subtypes, both in terms of cancer cell ligands interacting with CAF receptors and CAF ligands engaging cancer cell receptors. In contrast, healthy and pre-cancerous epithelial cells showed fewer such interactions (Fig. 2g,h). To assess cancer-CAF interactions associated with patient survival, we analyzed the expression of all ligand and receptor genes in PDAC cancer and CAF populations. We identified key microenvironment components, including CTHRC1, THBS2, and EFEMP1, which showed both a high percentage of patient expression and significant correlation with poor survival outcomes. Similarly, key receptors such as TSPAN1, ITGB4, ITGA2, and SDC1 were highly expressed across patients and also correlated with poor survival (Fig. 2i,j). We next assessed ligand-receptor interactions specifically between cancer and CAF subtypes. FN1, COL1A1, and THBS1 emerged as the most frequently expressed ligands in CAFs, interacting with receptors such as SDC1, ITGA2, and ITGA3 across multiple CAF-cancer cell type pairs. Conversely, ITGB1 and LRP1 were the most commonly expressed ligands in cancer cells, engaging receptors such as ADAM9, SERPIN, and LAMC2 in numerous cancer-CAF cell type interactions (Extended Data Fig. 3f). Taken together the integrated atlas identifies PDAC cellular expression states that are highly prevalent across most patients, and provides a substantial reference cohort for benchmarking in vitro models.

### Establishment of a cancer-CAF tumoroid culture system

We developed a tumoroid system to investigate cancer-CAF interactions (Extended Data Table 3). We first generated cancer cyst organoids (CCOs) from primary biopsies of PDAC patients and maintained CCOs in long-term culture following previously established protocols (Boj et al., 2015). scRNA-seq revealed that heterogeneity in cell cycle states was a major source of transcriptional variation within CCO cultures (Extended Data Fig. 3a-g). Comparison of CCO scRNA-seq profiles with the primary cancer cell atlas showed that CCO-derived cancer cells exhibited high transcriptional similarity to classical cancer cell subtypes (Extended Data Fig. 3g). We used CCOs to establish a stroma-rich PDAC tumoroid co-culture system by embedding cancer cells and CAFs in a three-dimensional matrix (Fig. 3a). Notably, the CAFs were not patient-matched, and the same CAF line was used as a base line throughout the study. Over the course of 14 days, we observed the emergence of complex cellular architecture within the tumoroids in which cancer cells formed ductal structures and CAFs produced an abundant extracellular matrix composed of vimentin, fibronectin, and collagen fibers forming a meshwork around the cancer cells (Fig. 3b).

**Figure 3.**
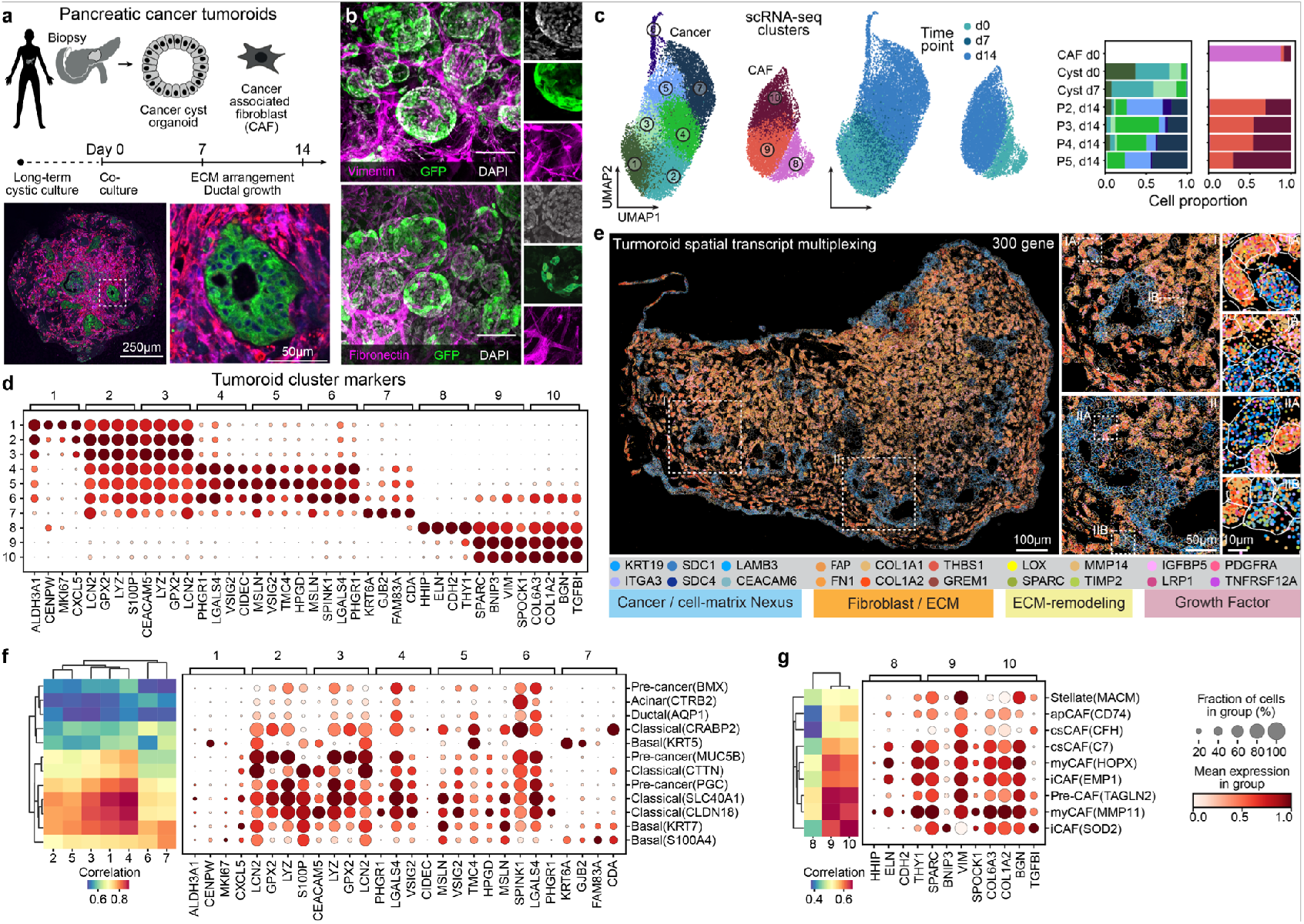
Modular tumoroids model pancreatic cancer cell interactions. (a) Long-term CCO cultures derived from patients with PDAC were co-cultured in a 3D matrix with CAFs, which self-organized into a complex tumoroid microenvironment. Over 14 days, fibrous connective tissue formed, vessels sprouted and organized, and cancer cells developed into 3D glandular structures within multilineage tumoroids. (b) Whole-mount immunohistochemistry on optically cleared tumoroids showing staining for Vimentin (pink, D) and Fibronectin (pink, E). Cancer cells express EGFP, and nuclei are labeled with DAPI (white). Scale bar: 100 µm. (c) scRNA-seq was performed on input cells from 2D CAF mono-cultures at Day 0 and Day 7, and from co-cultured tumoroids at Day 14. UMAP embedding of the scRNA-seq data is shown, colored by Leiden cluster (left) and by time point (right). The bar plot summarizes the proportion of each cell cluster across all samples. (d) Dotplot showing the normalized expression of top marker genes for each identified cluster. (e) MERSCOPE spatial transcriptomics in tumoroids. Four representative gene transcript clusters were detected in a tumoroid and visualized in a zoomed-in view. (f) Heatmap showing the correlation between cancer cell clusters identified by scRNA-seq in tumoroids and primary cancer cell types from the atlas in Fig. 2b. Right: dotplot showing expression of marker genes for each cancer cluster in Fig. 2d across the primary cancer atlas cell types. (g) Heatmap showing the correlation between CAF clusters in tumoroids and primary CAF subtypes from the atlas in Fig. 2c. Right: dotplot showing marker gene expression for each CAF cluster in Fig. 2d across primary CAF subtypes.

We performed scRNA-seq to assess cellular heterogeneity within the tumoroids from 4 patients at day 14 of co-culture and compared these profiles with those from CCO and CAF mono-cultures (Fig. 3c). The analysis revealed 10 distinct clusters, representing both cancer cell (C1-C7) and CAF states (C8-C10), with representative marker genes identified for each cluster (Fig. 3d). Cancer cells from CCOs were predominantly enriched in C1-C3, whereas cancer cells from tumoroids were enriched in C4-C7. Similarly, CAFs from mono-culture primarily localized to C8, while CAFs from tumoroids were enriched in C9 and C10. Tumoroid CAF C9 and C10 induced a consortium of extracellular matrix proteins (COL1A2, COL6A3) and genes associated with inflammatory response, transforming growth factor beta signaling, and hypoxia response, such as TGFB1, compared to C8. Marker gene analysis of the scRNA-seq clusters revealed that adenocarcinoma-associated genes such as GPX2, LYZ, S100P, and CEACAM5 were expressed across all cancer cell clusters. In contrast, some PDAC-specific genes including LGALS4, SPINK1, and MSLN were predominantly expressed in cancer cell clusters derived from tumoroids. Notably, C7 showed a distinct transcriptional profile, marked by the expression of basal-like cancer cell markers such as KRT6A and GJB2, suggesting a divergent or more aggressive cancer cell state emerges within the tumoroid (Fig. 3d). These data show that the multilineage tumoroid microenvironment induces morphological and molecular cell state changes across different cell types.

### Spatial analysis reveals interacting tumoroid states

To investigate the spatial organization of gene expression, we applied Multiplexed Error-Robust Fluorescence In Situ Hybridization (MERFISH (Chen et al., 2015), adapted by MERSCOPE) to tumoroid sections (Fig. 3e; Extended Data Fig. 5a-c; Extended Data Table 3). Following cell segmentation, we obtained a total of 69,870 cells from 27 tumoroids, with three tissue slices analyzed per tumoroid. UMAP embedding of the spatial transcriptomic data identified seven transcriptionally distinct clusters, comprising three cancer cell clusters (C1-C3) and four CAF clusters (C4-C7) that occurred in similar proportions across tumoroid sections (Extended Data Fig. 5d,e). Neighborhood enrichment analysis revealed spatial co-localization patterns among specific clusters, indicating organized cell-cell interactions within the tumoroid microenvironment (Extended Data Fig. 5f,g). Notably, cancer cell clusters C1 and C2 preferentially localized near CAF cluster C4 in the tumoroid interior, while cancer cluster C3 was spatially adjacent to CAF clusters C5-C7 towards the periphery. Differential gene expression analysis between clusters C2 vs. C3 and C4 vs. C5/6/7 identified key marker genes that distinguish epithelial-like (cancer) and mesenchymal-like (CAF) regions (Extended Data Fig. 5h,i). Mapping the top differentially expressed genes to previously defined cell states from scRNA-seq revealed that spatial cluster C2 aligned more closely with cancer clusters C5-C7, whereas C3 resembled cancer clusters C1-C4 (Extended Data Fig. 5j). We did not identify a clear spatial correspondence to the previously identified CAF clusters from scRNA-seq, potentially due to the restricted number of genes profiled in the MERSCOPE experiment. Overall, this spatial investigation resolved cell states, localizing adjacent cancer cell interactions with the CAF-induced extracellular microenvironment.

### Reference atlas comparison to assess tumoroid cell states

To benchmark tumoroid cell states, we compared the tumoroid scRNA-seq data with the PDAC cell atlas. Pseudobulk analysis followed by correlation between PDAC epithelial states and tumoroid cancer clusters revealed two major transcriptional groups within the tumoroid cancer cells (Fig. 3f). Clusters C1 to C5 showed high similarity to classical-like cancer cells in PDAC tissue, whereas clusters C6 and C7 were more closely aligned with basal-like cancer cells (Fig. 3f). The gene expression profiles of tumoroid clusters also mirrored those observed in PDAC primary epithelial cells (Fig. 3f). For example, C2 and C3 feature genes such as LCN2, GPX2, LYZ, SP100P, and CEACAM5, which are also highly expressed in classical-like cancer cells in the PDAC atlas. In contrast, C7 expresses feature genes including KRT6A, GJB2, and FAM83A, which are enriched in basal-like cancer cells in the atlas. Since C1-C5 cells originate from early-stage time points and C6-C7 emerge at later stages, these findings suggest that the tumoroid model recapitulates a classical-like transcriptional state early in culture and progressively shifts toward a basal-like state over time. A similar analysis was performed for CAF cells (Fig. 3g). Correlation and feature gene expression analyses showed that C8 cells, primarily from day 0, exhibited low similarity to primary tissue CAF states, whereas C9 and C10 cells correlated strongly with both myCAF and iCAF states in the PDAC atlas (Fig. 3g). Altogether, these findings indicate that the tumoroid model can capture the cell stages observed in the PDAC primary tissue atlas and that both cancer and CAF cells in later-stage tumoroids more closely resemble primary tissue states, highlighting the ability of the model to recapitulate in vivo-like cell states over time.

We next assessed the reproducibility of tumoroid CAF states through comparison to the integrated atlas cohort. We generated cancer cyst and CAF lines from the same biopsy of an additional individual (patient 7), and used these to establish an autologous CAF/cancer stromal tumoroid (Extended Data Fig. 6a). scRNA-seq profiling was performed for mono-cultured CAFs and CCOs exposed to growth factor supplemented media (day 0), CCOs in minimal media (day 7 and day 14) and tumoroids (day 7 and day 14) to assess heterogeneity and consistency of tumoroid states (Extended Data Fig. 6b). After integration, the cells grouped into 7 clusters representing mono-cultured CAF (C1), tumoroid CAF (C2,3), CCO cancer (C4,5,6), tumoroid cancer cells (C6,7) (Extended Data Fig. 6c,d). We observed differences in gene expression between cancer cells in CCOs and tumoroids, with tumoroid cancer cell states possessing similar signatures as previously detected (Extended Data Fig. 6e,f). Tumoroid CAF clusters from this patient presented a mixture of myCAF and iCAF features, which were particularly prominent compared to 2D culture conditions (Extended Data Fig. 6f). These data indicate relevant variation across patient lines, however cancer cells temporally converge to similar transcriptome states within the tumoroid co-culture.

### Modular incorporation of endothelial cells into tumoroids

Endothelial cells (ECs) can be co-cultured with cancer cells and CAFs, with progressive organization and formation of ductal structures, ECM, and vessel networks within the tumoroid by day 14 (Extended Data Fig. 7a-c). Single-cell RNA-seq revealed a maturation trajectory, with an early EC cluster (C6) shifting toward a more differentiated state (C7) marked by high VWF and CDH5 expression (Extended Data Fig. 7d-h). Cancer cells changed over time, with scRNA-seq revealing that PDAC-associated genes, including LCN2, AGR2, TSPAN8, and CEACAM5 were expressed across clusters with distinct dynamics, while cancer cells in CAF tumoroids showed enrichment of genes involved in metabolic homeostasis. LCN2 was identified as an early-induced gene (C9), and may serve as a potential early biomarker of cancer metabolic responses to CAF signaling. Basel-like cancer markers GJB2 and EGLN3 were more strongly enriched at later stages (C8), particularly in CAF-containing tumoroids, indicating that prolonged stromal interactions may promote relevant states.

Fibroblast identity determined the trajectories of tumoroid development. Normal fibroblasts (NFs) derived from the histologically normal portion of PDAC tissue supported less developed vascular networks and smaller ductal structures within tumoroids, and shifted gene expression by inducing focal adhesion-related genes such as AKAP12 at early stages (Extended Data Fig. 8a-f). NFs also enriched tumor-associated genes including SPON2 and MEG3 (C1) (Extended Data Fig. 8e,f). In contrast, CAFs continuously remodeled the extracellular matrix by inducing COL1A1, COL1A2, COL3A1, and COL6A3, while reinforcing PDAC signature genes (LCN2, TFF3, AGR2, TSPAN8) over 14 days, resulting in robust vessel development and larger, more complex ductal structures (Extended Data Fig. 8e-f). We validated low expression of AKAP12 and CAF-specific induction of TFF3 and LCN2 by immunohistochemistry (Extended Data Fig. 8g-i). LCN2 was also highly expressed in tumor regions of primary PDAC compared to normal pancreas (Extended Data Fig. 8j). Integrative single-cell analysis across multiple patients further showed consistent enrichment of CAF-associated genes in tumoroid fibroblast clusters (Extended Data Fig. 9a-f). Altogether, these data indicate that the system develops through sequential stages, where CAFs drive progressive and sustained remodeling generating vascularized, stroma-rich, and transcriptionally aggressive tumoroids resembling primary PDAC lesions.

### Interaction analysis and tumoroid perturbation identifies SDC1 as a PDAC central regulator

We identified many prevalent CAF-cancer interactions observed in primary tissues that were preserved in the tumoroid system (Fig. 4a, Extended Data Fig. 10a,b). Consistent with findings from primary PDAC samples, SDC1 emerged as a major receptor on cancer cells, capable of receiving signals from multiple CAF-derived ligands (Fig. 4a-c,, Extended Data Fig. 10a,b). These ligands include THBS1, FGF2, TNC, FN1, and collagen proteins, consistent with the known role of SDC1 as a multifunctional adhesion receptor and modulator of microenvironmental signaling (Yuan et al., 2023; Sleeboom et al., 2024). Previous studies have shown SDC1 to be upregulated via KRAS-driven recycling and essential for macropinocytosis in pancreatic cancer (Yao et al., 2019; Theardy et al., 2025). SDC1 engagement in extracellular matrix remodeling and ligand binding has been implicated in promoting tumor progression, proliferation, and metastasis, and is associated with poor patient prognosis (Ni et al., 2017; Chen et al., 2020; Topalovski and Brekken, 2016; Jacquemin et al., 2020; Bray et al., 2019; Nagpal et al., 2017). In line with these associations, Kaplan-Meier analysis of PDAC patients revealed significantly worse outcomes in cases with high SDC1 expression (Extended Data Fig. 10c). Immunofluorescence analysis revealed increased SDC1 protein levels at day 14 compared to day 7 tumoroids, predominantly localized on the cancer cell membrane (Fig. 4d). This spatial pattern was also validated in primary PDAC resections (Fig. 4e).

**Figure 4.**
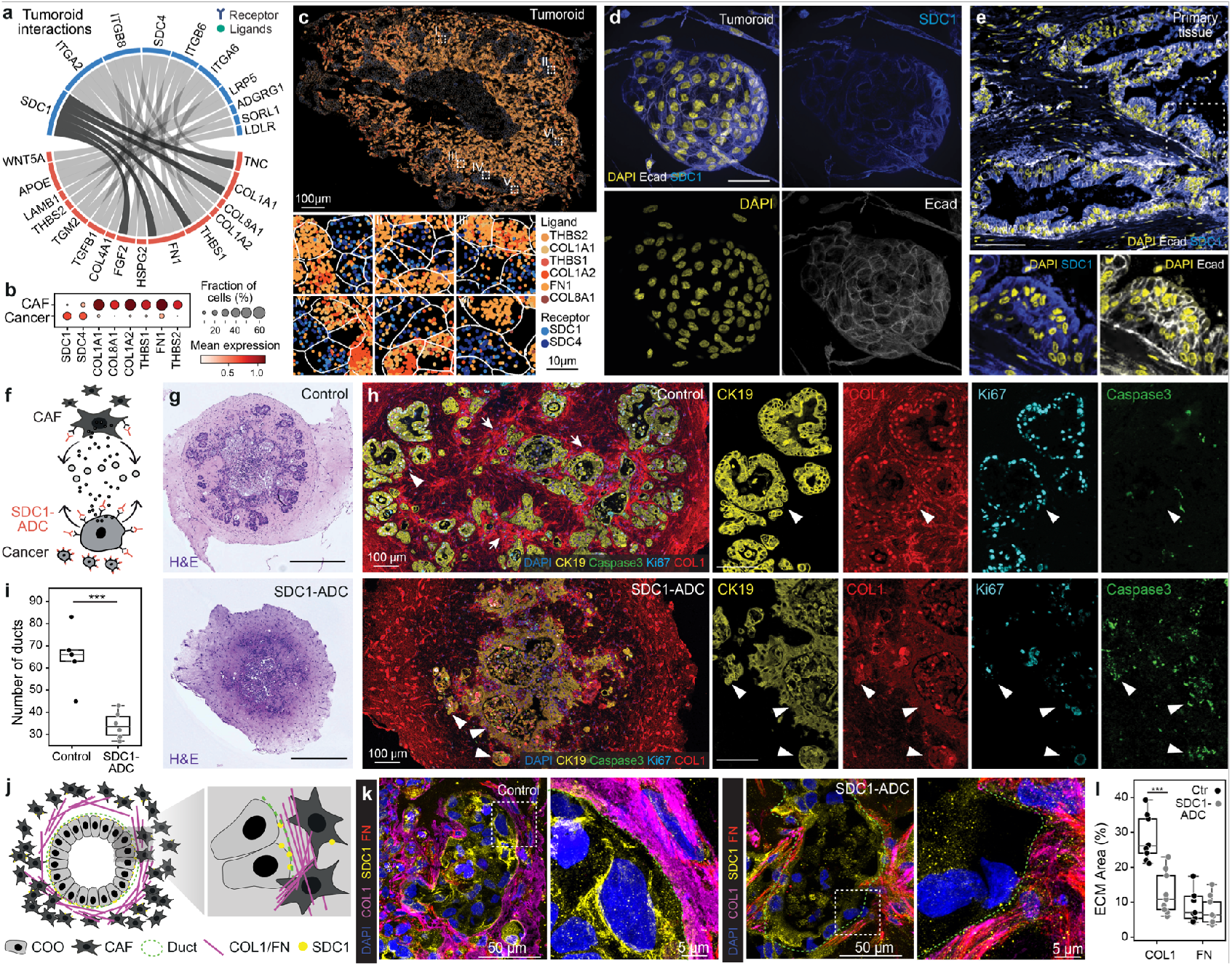
SDC1 is a therapeutic target regulating tumoroid growth. (a) Ribbon plots illustrating ligand-receptor (LR) interactions from CAFs to cancer cells in CAF-tumoroids. (b) Dotplot showing the expression of selected ligand and receptor genes in the tumoroid scRNA-seq data. (c) Spatial transcriptomics of selected ligand and receptor gene expression in tumoroids using MERSCOPE. (d) Immunofluorescence staining of pancreatic tumoroids at day 14, showing SDC1 (blue), E-cadherin (Ecad, gray), and DAPI (nuclei, yellow). Scale bar: 50 µm. (e) Immunofluorescence staining of primary human pancreatic cancer tissue showing expression of SDC1 (blue), E-cadherin (gray), and DAPI (yellow). Scale bar: 100 µm. (f) Schematic representation of anti-SDC1 antibody-drug conjugate (SDC1-ADC) targeting in the tumoroid model. (g) Representative images of tumoroids treated with control or SDC1-ADC, stained with H&E, and immunofluorescently labeled for DAPI (blue), KI67 (cyan), CK19 (yellow), cleaved caspase-3 (green), and COL1 (red). (h) Duct-like structures in control (upper) vs. SDC1-ADC-treated (lower) tumoroids, with CK19, Caspase-3, Ki67, and COL1 immunofluorescence staining shown on the right. (i) Boxplot illustrating the number of ducts in control and SDC1-ADC-treated tumoroids (^*^p < 0.05, ^**^p < 0.01, ^***^p < 0.001). (j) Schematic representation of super-resolution microscopy imaging and analysis in tumoroids. (k) Super-resolution microscopy imaging of tumoroids in control (upper) vs. SDC1-ADC-treated (lower) conditions, with a zoomed-in image on the right. (l) Quantification of ECM area percentages for COL1 and FN in control and SDC1-ADC-treated tumoroids (^*^p < 0.05, ^**^p < 0.01, ^*^p < 0.001).

We performed functional perturbation of SDC1 using an antibody-drug conjugate (SDC1-ADC) carrying the cytotoxic payload DM4 (Fig. 4f). Brightfield imaging and organoid formation assays demonstrated selective sensitivity of PDAC organoids to SDC1-ADC compared to healthy organoids (Extended Data Fig. 10d,e). High-resolution confocal imaging revealed that SDC1-ADC treatment to tumoroids led to collapse of duct-like structures, with loss of polarized epithelial organization and reduced outgrowth compared to untreated controls (Fig. 4g-i). Multiplex immunofluorescence showed increased apoptosis (cleaved Caspase-3) and reduced proliferation (Ki-67), consistent with impaired cancer cell survival (Fig.4h). Because SDC1 integrates signals from multiple CAF-derived ligands, we next assessed how its blockade affected cancer-CAF-ECM interactions. Super-resolution AiryScan microscopy revealed marked disruption of this interaction axis, with COL1 and FN redistributed into thinner fibrils surrounding smaller pericellular regions, and with THBS1 and TNC also affected (Fig. 4j-l; Extended Data Fig. 10f,g). These results demonstrate that SDC1 is essential for cancer cell growth and is involved in coordinating CAF-dependent ECM remodeling, underscoring its central role in maintaining tumor-stroma communication in PDAC. Together, these findings highlight SDC1 as a central node in tumor-stroma communication and demonstrate that multilineage PDAC tumoroids provide a powerful platform to dissect cancer-CAF interactions with therapeutic potential.

## Discussion

We demonstrate a strategy to dissect PDAC by combining an integrated single-cell transcriptomic atlas with a modular, stroma-rich tumoroid model. The PDAC atlas, built from over 670,000 cells across 200 samples, defines conserved cancer and stromal cell subtypes and reveals shared gene programs and ligand-receptor interactions across patients. This atlas provides an important reference for identifying universal PDAC signatures and variable features, and evaluating the fidelity of in vitro models. Using this atlas as a benchmark, we show that co-cultured tumoroids derived from patient cancer cells and CAFs recapitulate key features of the primary tumor microenvironment. Single-cell and spatial transcriptomic profiling confirmed the presence of major epithelial and mesenchymal cell states observed in the atlas, including classical and basal-like cancer subtypes and multiple CAF phenotypes. These cellular states exhibit dynamic behavior within the tumoroid context, including progressive remodeling, matrix deposition, and cancer cell migration. CAFs modulate microenvironmental interactions and exert strong phenotypic influence on cancer cells, notably via the receptor SDC1. Blocking SDC1 impairs ductal growth within tumoroids, highlighting a potentially targetable mechanism of CAF-driven cancer progression. Moreover, cell states associated with poor prognosis in primary tumors, such as migratory basal-like cancer cells and matrix-producing myofibroblastic CAFs, co-emerge within tumoroids, underscoring the clinical relevance of this model. While this work focuses primarily on cancer-fibroblast interactions, we also highlight the modularity of the system and how fibroblasts support endothelial cell incorporation and organization within the tumoroid. Future studies can harness this modularity to explore how cancer, mesenchymal, endothelial, immune, neural and other cell types interact to coordinate PDAC intra-tumoral complexity. The combination of a reference atlas with tractable co-culture models offer new inroads for functional studies and translational discovery in pancreatic cancer (Okuda et al., 2025).

## Author contributions

QX integrated the PDAC atlas and conducted atlas analyses. RO, CE established and banked the organoids and fibroblasts lines from biopsy tissues with support SP. RO developed the tumoroid protocol with support from YU and HT. RO generated tumoroids and performed IHC experiments. RO, QX, and BG performed tumoroid single-cell RNA-seq experiments and analyzed the data. MS performed the MERSCOPE experiment. QX and CH analyzed the MERSCOPE data. ML performed the super-resolution imaging experiment. Patient material was provided by CE, KJ, SP, SM, YM, TY, and HT. QX, RO, BG, BT, and JGC designed the study, interpreted the results, and wrote the manuscript. All authors discussed the results and approved the final version of the manuscript.

## Acknowledgements

We thank the Camp and Treutlein labs, and Keisuke Sekine for helpful discussions. We extend our gratitude to the UCSC cell browser team (https://cells.ucsc.edu/) for their invaluable assistance in organizing and facilitating the publication of our atlas as a cell browser. We are grateful to Annegret Fischer (pRED Data&Analytics) for computational support with atlas dataset procurement. The results shown here are in part based upon data generated by the TCGA Research Network: https://www.cancer.gov/tcga. This work was supported in part by Chan Zuckerberg Initiative DAF, an advised fund of the Silicon Valley Community Foundation CZF2019-002440 (J.G.C., B.T.), the Swiss National Science Foundation (Project Grant-310030_84795, J.G.C.; Project Grant-310030_192604, B.T.), and the National Center of Competence in Research Molecular Systems Engineering (B.T.). R.O. was supported by the Japanese Society for the Promotion of Science (JSPS).

## Data availability

The PDAC atlas, along with sub-atlases of epithelial and mesenchymal cells (including raw and normalized counts, integrated embeddings, cell type annotations, and technical metadata), is publicly available via the UCSC Cell Browser at https://cells.ucsc.edu/?ds=pdac-atlas. The GRCh38 genome assembly used in this study is accessible at https://www.ncbi.nlm.nih.gov/datasets/genome/GCF_-000001405.26/. Single-cell RNA-seq data generated from all PDAC tumoroid samples in this study will be deposited in the Gene Expression Omnibus (GEO) database.

## Code availability and analytic reproducibility

The code for scRNA-seq cell type annotation is available as a Python package maintained on GitHub at https://github.com/devsystemslab/snapseed. All analysis scripts used in this study are available at https://github.com/devsystemslab/pdac_atlas.

## Supplemental information (5)

Extended Data Table 1: Sample information included in the PDAC atlas.

Extended Data Table 2: Differentially expressed genes in cancer and CAF cells detected in the PDAC atlas.

Extended Data Table 3: Patient information utilized in this study.

Extended Data Table 4: Gene list used in the MER-SCOPE experiment.

Extended Data Table 5: Primary and secondary anti-bodies used in this study.

## Methods

### PDAC single cell atlas data collection

The scRNA-seq raw FASTQ files or count data used in this study were obtained from their respective original publications. When raw FASTQ files were unavailable, reads were aligned to the GRCh38 genome with Ensembl 98 gene annotation using STARsolo (Kaminow et al., 2021). In cases where raw FASTQ files could not be accessed, we instead downloaded preprocessed raw count matrices. These downloaded counts were then merged with those obtained from the realigned reads for downstream analysis.

### Data Integration Benchmarking

To benchmark single-cell RNA-seq data integration methods in PDAC datasets, 12 methods were evaluated, including PCA, Seurat (v3, v4, and v5), scVI, scANVI, scPoli, bbknn, harmony, combat, CSS (Pearson), and CSS (Spearman) (Polański et al., 2020; Xu et al., 2021; Lopez et al., 2018; Korsunsky et al., 2019; He et al., 2020; Büttner et al., 2019; De Donno et al., 2023; Hao et al., 2024). The scIB benchmarking tool (Luecken et al., 2022) was employed to evaluate and compare the performance of these methods in integrating the samples. Due to the computational cost associated with Seurat methods, integration was performed on a randomly selected subset of 25 samples for Seurat-based approaches.

### PDAC atlas integration

PDAC atlas data were processed using Scanpy (Wolf et al., 2018). To integrate the data, count matrices from all samples were combined into a unified dataset. For downstream analysis, only protein-coding and long non-coding RNA (lncRNA) genes were retained, while low-quality cells were filtered out. Raw counts were normalized to 10,000 total counts per cell and log-transformed. Highly variable genes (HVGs) were identified using Scanpy’s default settings, selecting the top 3,000 HVGs for further analysis.

Cell type annotation was performed separately for each sample using the Snapseed method (https://github.com/devsystemslab/snapseed). Principal component analysis (PCA) was applied to the normalized data, and the top 30 principal components (PCs) were used to construct a K-nearest neighbors (KNN) graph. Cell clustering was performed using the Leiden algorithm with a resolution of 2, enabling the identification of distinct cellular populations based on gene expression profiles. Annotation was guided by previously established marker genes, as follows:

Epithelial cells (KRT8, EPCAM, REG4, KRT5, CLDN18) include acinar (CTRB2, CPB1), ductal cells (AQP1, AMBP), classical (GCNT3, REG4, CLDN18), and basal-like (KRT5, KRT6A). Mesenchymal cells (DCN, PDGFRB) include stellate/pericyte-like cells (FABP4, MCAM), myofibroblastic CAFs (MMP11), inflammatory CAFs (IL6, LIF, CXCL1, CXCL2), and antigen-presenting CAFs (HLA-DRA, HLA-DRB1, CD74). Lymphoid cells (CD3E, CD3D, NKG7, KLRK1, CD79A, MS4A1) include CCR7, IL7R, FOXP3, and CXCL13 CD4 T cells; GZMH, GZMK, and ITGA1 CD8 T cells; ISG15 T cells; GNLY and XCL1 NK cells; proliferating T/NK cells (MKI67); and B/plasma cells (CD79A, MS4A1). Myeloid cells (CD68, CD14) include SPP1 (SPP1, MARCO) and C1QC macrophages (C1QA, C1QB, C1QC); MDSCs (S100A8, S100A9, S100A12); monocytes (FCGR3A, CDKN1C); cDC1 (CLEC9A, XCR1), cDC2 (CD1C, FCER1A), and cDC3 (CCL19, CCL22, CCR7); pDC (LILRA4, PLD4); and proliferating myeloid cells (MKI67). Endothelial cells (VWF, CDH5, KDR) consist of vascular (FLT1) and lymphatic (FLT4) subtypes. Endocrine cells (ING, GCG, IAPP, CHGA, SST, PPY, NKX6-1, HEPACAM2) include alpha (GCG), beta (IAPP, NKX6-1, HEPACAM2), delta (SST), PP (PPY), and epsilon (GHRL) cells. Additional cell types identified include mast cells (KIT), erythrocytes (ALAS2, SLC4A1, SPTA1, KLF1), platelets (ITGB3, ITGA2B, GP1BA), and Schwann cells (CDH19).

Cell types were assigned within each cluster based on these marker genes to enhance annotation accuracy. After annotation, cells from all samples were merged into a unified dataset, and the top 3,000 highly variable genes were selected for integration. All cells were integrated using the scPoli method (De Donno et al., 2023), with cell_type as the integration key. The following scPoli parameters were used: early_stopping_metric = val_prototype_loss, mode = min, threshold = 0, patience = 20, reduce_lr = enabled (lr_patience = 13, lr_factor = 0.1), n_epochs = 12, pretraining_epochs = 10, eta = 10, and alpha_epoch_anneal = 100.

### Hierarchical clustering of cell proportions with ordinal metadata

To improve the interpretability and biological relevance of hierarchical clustering based on cell type proportions, we developed a method that integrates scRNA-seq cell proportions with external ordinal metadata (e.g., clinical stage). Instead of clustering solely based on cell type proportions, our approach incorporates both compositional and metadata-derived distances at the distance matrix level prior to linkage-based clustering. Let *X* ∈ *R*^*n*×*p*^ be the cell proportions matrix with n samples and p features. Let *m* ∈{*l*_1_, *l*_2_, …, *l*_*k*_}^*n*^ be a list of ordinal metadata labels (e.g., clinical stage I, II, III, possibly including unknown). Each label is mapped to a numeric value using a predefined level map *f* : *l*_*i*_ →*R*. Missing or unknown values are imputed using the median, mean, or a fixed value as specified. Pairwise Euclidean distances are computed separately on the expression matrix *D*_*X*_ = *pdist*(*X*) and the numeric metadata vector *D*_*m*_ = *pdist*(*m*). These two distance matrices are combined via a weighted linear sum: *D*_*c*_*ombined* = *aD*_*X*_ + *bD*_*m*_, where α and β are user-defined weights (default: α=0.8, β=0.2). Hierarchical clustering is then performed on *D*_*c*_*ombined* using Ward’s method. To enhance the visual interpretability of dendrograms and associated heatmaps, we applied the optimal_leaf_ordering function from the SciPy package (Virtanen et al., 2020) to reorder dendrogram leaves.

### Differential gene expression analysis in PDAC

For epithelial cells, acinar and ductal cells were grouped as normal cells, while all other epithelial subtypes were grouped as cancer cells. For mesenchymal cells, stellate cells were considered normal, and all other mesenchymal subtypes were grouped as CAFs. For each patient, the average (bulk) expression of genes in normal and cancer cell groups was calculated. If a given cell group contained fewer than 10 cells, additional cells of the same type were randomly sampled from other patients to reach a minimum of 10 cells per group. Differential gene expression across patients was assessed using the paired Wilcoxon rank-sum test for all genes. A prioritization score (PI score) was defined as the product of the log fold change and the negative log p-value (*PI* = *logFC* × − *log*_10_(*p*)). Genes with a PI score > 1 were considered differentially expressed across patients.

### Pathway enrichment analysis

Pathway enrichment analysis was performed using the Enrichr tool implemented in the gseapy package (Fang et al., 2023). Differentially expressed genes (DEGs) from cancer cells, CAFs, or their overlap were used as input gene sets. Enrichment was tested against the curated WikiPathways 2024 database (https://www.wikipathways.org/). Significantly enriched pathways were identified based on adjusted p-values, with a cutoff of 0.5 applied to filter results. Enrichment significance was transformed by calculating the negative logarithm of the adjusted p-values (- log10 padj) for visualization and ranking. The results were then sorted in descending order of significance to prioritize the most relevant pathways.

### Survival analysis

To investigate the association between gene expression and patient prognosis in PDAC, we performed gene-wise survival analysis using RNA-seq and clinical data from the TCGA cohort, accessed via the RTCGA package. For each gene, normalized expression values were retrieved by matching gene symbols. Clinical records were filtered to retain samples with available survival time and status. Overall survival was defined as the number of days from diagnosis to death or last follow-up, with deceased patients coded as 1 and censored patients as 0. Expression and clinical data were merged using patient identifiers. Patients were stratified into “High” and “Low” expression groups based on the 25th percentile of expression as a threshold. Kaplan-Meier survival curves were generated using the survfit function from the survival package, and survival differences between groups were evaluated using the log-rank test via the surv_pvalue function from the survminer package. To control for directionality, we defined the “worst” group as the one with the smallest area under the survival curve (AUC), computed from the Kaplan-Meier estimates. If the “Low” expression group was associated with poorer prognosis, the corresponding p-value was reassigned to 1, thereby excluding genes where high expression was not linked to worse outcomes.

### Ligand receptor analysis

To investigate cell-cell communication in the PDAC atlas and tumoroid model, ligand-receptor (LR) interaction analysis was performed. All curated LR pairs from the Lignature method (Xin et al., 2025) were downloaded for downstream analysis. For the PDAC atlas (level 1 cell types), cell-type-specific differentially expressed genes (DEGs) were identified using the Wilcoxon rank-sum test across the entire atlas. For each cell type, the top 500 DEGs were selected based on p-value. LR pairs were defined when both the ligand and receptor genes were among the DEGs in the corresponding sender-receiver cell type pairs. Identified interactions were summarized in a heatmap to visualize intercellular communication. For each cancer or CAF sub-atlas, DEGs were similarly identified using the Wilcoxon rank-sum test. Genes with p-values < 0.01 were considered significant and ranked by p-value. If more than 500 significant genes were found, the top 500 were retained. LR pairs were defined based on the co-expression of ligands and receptors among DEGs from the relevant CAF-cancer cell type pairs. For the tumoroid model, all cancer cells and all CAFs were merged, and DEGs were identified separately for each population. The top 500 DEGs for cancer and CAF cells were selected, and LR pairs were identified where both genes were among the DEGs of the CAF-cancer cell pair.

### Establishment of cystic organoid and fibroblast cultures

Human PDAC tissue samples and associated data were obtained (Extended Data Table 3). Organoids and stromal cells were established from clinical specimens obtained and processed at the University of Basel (Basel, Switzerland) and the Kanagawa Cancer Center (Yokohama, Japan). Informed consent was obtained from all patients prior to sample collection, and protocols were approved by the respective institutional ethics committees. Tumor and adjacent healthy tissues were collected by surgical resection. The cancer cyst organoid (CCO) culture method from PDAC tumor specimens is briefly described below (Boj et al., 2015). The surgical tissue is washed several times with Dulbecco’s phosphate buffered saline (DPBS). The tissue was finely chopped using surgical scissors and a scalpel. The tissue was transferred to a 50 ml tube and washed again with DPBS. The washed tissue was digested with LiberaseTM (Roche) at 37°C for 40-60 minutes. Tissues were enzymatically treated and then washed with DMEM containing 10% fetal bovine serum (FBS, Sigma) to stop the enzymatic reaction. The obtained pancreatic cancer cells were embedded in growth factor reduced (GFR) Matrigel (Corning) and cultured in the following complete medium. DMEM/F12(Thermo), Primocin (1mg/ml, InvivoGen), GlutaMAX (1x, Invitrogen), 1x B27(1x, Invitrogen), Gastrin, N-acetyl-L-cysteine (1mM, Sigma), Nicotinamide (10mM, Sigma), A83-01(Tocris, 0.5uM), Noggin (Peprotech, 0.1ug/ml), R-Spondin1 (Peprotech, 100ng/ml), Wnt3A(R&D, 50ng/ml), EGF (Peprotech, 50 ng/ml), FGF10 (Peprotech, 100ng/ml). Y-27632 (Sigma, 10uM) was added for only one day after starting the organoid culture, and on the following day, the cells were cultured in a complete medium without Y-27632. The medium was changed 2 to 3 times a week. For establishment of fibroblasts, healthy pancreatic tissue and cancer tissue were treated with Liberase and the collected cells were washed with DPBS several times. Subsequently, cells were suspended in Mesenchymal stem cell growth media (MSCGM, Lonza) and seeded on a culture plate. The media was changed 2-3 times a week. All cells were cultured under 5% CO2 in 20%O2 at 37°C. All lines used in the studies were verified as negative for mycoplasma before further experimentation.

### Tumoroid culture method

To establish a stroma-rich pancreatic tumoroid, pancreatic CCO cells, fibroblasts and human umbilical vein endothelial cells (HUVECs) were separately expanded and cultured. Pancreatic CCO cells were incubated with Triple EX (Gibco) for 7 minutes, and fibroblasts and HUVECs were incubated for 3 minutes at 37°C to generate a cellular suspension. To stop the enzymatic reaction by Triple EX, the cells were washed with DMEM/F12 medium containing 10% FBS and 1% Penicillin-Streptomycin (P/S, Gibco). The obtained cells were counted separately, and then 3 × 10^4^ cancer cells, 1.2 × 10^4^ HUVECs, 8 × 10^4^ fibroblasts were transferred to a tube coated with bovine serum albumin (1% BSA), mixed and centrifuged at 300 g. After removal of the supernatant, cell pellets were gently resuspended and 1.2 × 10^5^ cells were then seeded in 96 well plates coated with 50% Matrigel (Corning). The three types of cells made cell-cell interactions with each other and showed self organisation during the period of 24-48 hours. The reconstituted stromal-rich pancreatic tumoroid were cultured with 50% Endothelial Cell Growth Media (Lonza) and 50% DMEM/F12 medium. Culture mediums were exchanged every 24 hours. Tumoroid were cultured under 5% CO2 in 20%O2 at 37°C.

### Generation of reporter lines

For live imaging, HUVECs were infected with retroviruses expressing Kusabira-Orange (KO) and cancer cells were infected with a lentivirus expressing enhanced green fluorescent protein (EGFP) (Koike et al., 2004). Briefly, Human Embryonic Kidney (HEK) 283T cells were transfected with the retroviral vector pGCDNsam IRES-EGFP or KOFP (M. Onodera) for packaging at 293gag / pol (gp) and 293gpg (gp and VSV-G) to induce viral particle production. The culture supernatant of the retrovirus-producing cells was passed through a 0.45 mm filter (Whatman, GE Healthcare) and immediately used for infection. The firefly luciferase gene was subcloned into the CSII-EF-MCS-EGFP vector (RIKEN BRC) to generate the CSII-EF-Luc-IRES-EGFP construct. CSII-EF-Luc-IRES-EGFP plasmid and helper plasmid (293T cells were transfected with calcium phosphate using pCAG-HIVgp and pCMV-VSV-G-RSV-Rev, RIKEN BRC) to produce VSV-G pseudotyped lentivirus. The virus supernatant was recovered 46 hours after transfection, and filtered with a 0.45 µm filter. The virus supernatant was concentrated by ultracentrifugation.

### Whole-mount clearing imaging

Tumoroids were washed several times with PBS and fixed with 200ul of 4% (wt/vol) paraformaldehyde (PFA). Tumoroids were incubated on a horizontal shaker at 4ºC for 24 hours. PFA was then completely removed and fixed tumoroids were washed several times with PBT buffer (0.1% Tween (vol / vol)). Tumoroid washing buffer (TWB: 100ml of PBS with 0.2 g of BSA and 0.1% Triton X-100) was added to the wells and incubated on a horizontal shaker at 4ºC for 1 day to block tumoroid. The next day, the blocking reagent was completely removed from the well, and then 100 ul of TWB with primary antibodies was added to the wells and incubated on a horizontal shaker at 4ºC for 2 days. After immuno-labeling the tumoroid with the primary antibody, these reagents were removed from the wells, and then fresh TWB was added to the wells and was incubated on a horizontal shaker at 4ºC for 2 hours.

This process was performed 3 times to completely remove the antibodies from organoids. After the tumoroid were sufficiently washed with TWB, 100ul of TWB with secondary antibodies was added to the wells and incubated on a horizontal shaker at 4ºC for 1 day. After immuno-labeling the tumoroid with the secondary antibody, the secondary antibodies were removed from the wells and then fresh TWB was added to the wells and washed three times. Subsequently, 50ul of the fructose-glycerol clearing solution (Dekkers et al., 2019) was added to the well and incubated on a horizontal shaker at 4ºC, overnight. Cleared organoids were placed on a glass slide or in a glass-bottom plate and imaged on a spinning disc confocal microscope (Olympus SpinSR10 spinning disk confocal super resolution microscope, objective x10,x20,x30,x40,x60). Primary and secondary antibody information are described in Extended Data Table 5.

### FFPE processing and analysis of co-cultured tumoroid samples

Co-cultured tumoroid samples were washed three times with 1× DPBS and fixed with 4% paraformaldehyde (PFA) in a 96-well Clear TC-treated plate for 45 minutes at room temperature. After fixation, the wells were washed three more times, and the remaining liquid was aspirated. Preliquefied HistoGel (ThermoScientific) was distributed into biopsy cassettes, and samples were transferred from the plate into the HistoGel for embedding. Once polymerized, samples were dehydrated overnight in a vacuum filter processor (Sakura TissueTek VIP5) and embedded in liquid paraffin the following day. FFPE blocks were sectioned at a thickness of 3.5 µm using a microtome and mounted on Superfrost Plus Adhesion microscope slides (Epredia). Slides were then incubated overnight at 37 °C in a slide oven.

For H&E staining, slides were deparaffinized, rehydrated, and stained with hematoxylin (Sigma-Aldrich) for 5 minutes, followed by rinsing under running water for 5 minutes. Sections were differentiated in 0.3% acid alcohol for 20 seconds, washed again under running water for 5 minutes, and counterstained with eosin (Thermo Fisher) for 2 minutes. Slides were dehydrated, cleared, and mounted for imaging.

Multiplex immunofluorescence (mIF) staining was performed on FFPE slides using a Ventana Discovery Ultra automated tissue stainer (Roche Tissue Diagnostics). Slides were baked at 60 °C for 8 minutes, heated to 69 °C for deparaffinization in two cycles, and subjected to antigen retrieval with Tris-EDTA buffer (pH 7.8; Ventana) at 92 °C for 32 minutes. Blocking with Discovery Inhibitor (Ventana) for 16 minutes was followed by neutralization. Primary antibodies, diluted in Discovery Ab diluent (Ventana), were detected with HRP-conjugated secondary antibodies (OmniMap Ventana; Table S2), and Opal dyes (Akoya Biosciences) were applied. After each staining cycle, antibodies and HRP were neutralized and denatured, and subsequent cycles began with fresh blocking steps. Nuclei were counterstained with DAPI (Roche). Details of the antibodies are listed in Extended Data Table 5.

Stained slides were digitized using the Vectra Polaris multispectral imaging system (PerkinElmer) with MOTiF technology at 20× magnification, capturing seven fluorophores (Opal 480, 520, 570, 620, 690, 780, and DAPI). Slides were scanned in batches under the same settings to ensure consistency. Channel unmixing and image tiling were performed using PhenoChart (v1.0.12) and inForm (v2.4), and fused images were further analyzed using HALO (v3.2.1851.328) with HALO AI (Indica Labs).

### Super-resolution microscopy and SDC1-targeted treatment

Super-resolution AiryScan imaging was conducted using a Zeiss LSM980 confocal microscope with a Plan-Apochromat 63×/1.40 NA Oil DIC M27 objective. Imaging across four channels was performed in two experimental sets: Experiment 1 (COL1, FN, SDC1, and Hoechst) and Experiment 2 (THBS1, TNC, SDC1, and Hoechst). Laser power settings were optimized, with Experiment 1 using SBS LP 640 (1%), SBS LP 525 (0.6%), SBS LP 550 (1%), SBS LP 505 (0.4%) and Experiment 2 using SBS LP 640 (0.7%), SBS LP 525 (0.5%), SBS LP 550 (1%), SBS LP 505 (0.3%). AiryScan detection was operated in Super-Resolution mode, with z-stack acquisitions (total thickness: 9.88 µm, z-step size: 0.130 µm, xy pixel size: 0.035 µm).

Images were processed using the 3D AiryScan reconstruction algorithm in ZEN Blue software (Zeiss) and analyzed with a custom Python script. Binary masks were generated for each ligand channel via intensity thresholding. Mean fluorescence intensity was extracted and normalized (min-max scaling), while mask areas were expressed as the percentage of the field of view occupied by the ligand. Morphological features, such as fiber structures, were quantified using the maximum Feret diameter to assess ligand-associated fiber organization and remodeling between control and treated samples.

CAF-cancer cocultured tumoroids were treated with ADC-W-320 (Indatuximab Ravtansine, Anti-SDC1 (nBT062)-SPDB-DM4 ADC, Creative Biolabs) by replacing the medium on day 4 with 200 µL of coculture medium containing ADC at 5 µM. The medium was refreshed every 3 days, and the treatment continued until day 14. For cancer cyst and healthy pancreatic organoids, ADC-W-320 was applied at a concentration of 0.5, 1, or 5 µM starting on day 4 and maintained for 7 days (short-term) or 14 days (long-term), with medium changes every 3 days. Cultures were maintained at 37 °C in a 5% CO and 20% O environment and imaged at designated endpoints.

### Single-cell RNA-seq experiments

All samples were dissociated to single cells by specific enzymatic treatment. The cultured medium for stroma-rich tumoroid was removed from the wells and tumoroids washed three times with 1xDPBS. The tumoroids were collected in 5 ml tubes, after the DPBS was completely removed from the tube, TrypLE™ Select (Thermo) was added and incubated at 37ºC for 8 minutes (Gehart et al., 2019). After the incubation step, tumoroids were further dissociated by trituration. This incubation and trituration process was repeated 3 times to obtain a single cell suspension. The enzymatic dissociation was stopped by addition of cold BE-PBS (Cold PBS 1 ml with 0.04% BSA / (0.1 mM EDTA)) and remaining cellular clumps were removed by using 70um and 40um strainers. Fibroblasts and HUVECs were cultured on a 10 cm dish and dissociated to a single cell suspension using the same procedure as described above. Single cell suspensions were adjusted to an appropriate concentration to obtain approximately 2000-10000 cells per lane of a 10x Genomics microfluidic Chip G. Libraries were generated using 10x Genomics 3’ Gene Expression Kit (v3.1), following recommended protocol, and subsequently sequenced on NextSeq500, using 28-9-0-91 Read Configuration, as recommended by for Single Index libraries.

### Single-cell RNA-seq data preprocessing

CellRanger (v3.1.0, 10x Genomics) was used to extract unique molecular identifiers, cell barcodes, and genomic reads from the sequencing results of 10x Chromium experiments. Then, count matrices, including both protein coding and non-coding transcripts, were constructed aligning against the annotated human reference genome (GRCh38, v3.0.0, 10x Genomics). In order to remove potentially damaged or unhealthy cells and improve data quality, the following filtering steps were performed in addition to the built-in CellRanger filtering pipeline. Cells associated with over 20,000 transcripts, usually less than 1% of the total number of samples, were removed. Cells associated with a low number of transcripts (<1% of the total number of samples) were removed. Cells with over 15% of mitochondrial transcripts were removed. Transcripts mapping to ribosomal protein coding genes were ignored. Cells with <800 unique transcripts (<1% of the total number of samples) were removed together with transcripts detected in less than 10 samples.

### Normalization with Seurat

For normalization and variance stabilization of each scRNA-seq experiment’s molecular count data, we employed the modeling framework of SCTransform in Seurat v3 (Stuart et al., 2019). In brief, a model of technical noise in scRNA-seq data is computed using ‘regularized negative binomial regression’. The residuals for this model are normalized values that indicate divergence from the expected number of observed UMIs for a gene in a cell given the gene’s average expression in the population and cellular sequencing depth. Additionally, a curated list of cell cycle associated genes, available within Seurat, was used to estimate the contribution of cell cycle and remove this source of biological variation from each dataset in order to increase the signal deriving from more interesting processes. The residuals for the top 2,000 variable genes were used directly as input to computing the top 100 Principal Components (PCs) by PCA dimensionality reduction through the RunPCA() function in Seurat. Corrected UMI, which are converted from Pearson residuals and represent expected counts if all cells were sequenced at the same depth, were log-transformed and used for visualization and differential expression (DE) analysis.

### Doublet removal with DoubletFinder

For each scRNA-seq experiment DoubletFinder (McGinnis et al., 2019) was used to predict doublets in the sequencing data. In brief, this tool generates artificial doublets from existing scRNA-seq data by merging randomly selected cells which are then pre-processed together with real data and jointly embedded on a PCA space that serves as basis to find each cell’s proportion of artificial k nearest neighbors (pANN). For this step we restricted the dimension space to the top 50 PCs. Finally, pANN values are rank ordered according to the expected number of doublets and optimal cutoff is selected through ROC analysis across pN-pK parameter sweeps for each scRNA-seq dataset; pN describes the proportion of generated artificial doublets while pK defines the PC neighborhood size. In order to achieve maximal doublet prediction accuracy, mean-variance normalized bimodality coefficient (BCmvn) was leveraged. This provides a ground-truth-agnostic metric that coincides with pK values that maximize AUC in the data. To overcome DoubletFinder’s limited sensitivity to homotypic doublets, we consider doublet number estimates based on Poisson statistics with homotypic doublet proportion adjustment assuming 1/50,000 doublet formation rate the 10x Chromium droplet microfluidic cell loading.

### Tumoroid scRNA-seq integration

All scRNA-seq samples were processed and merged using the Scanpy framework. Raw counts were normalized to 10,000 total counts per cell and subsequently log-transformed. Highly variable genes (HVGs) were identified using Scanpy’s default settings, and the top 3,000 HVGs were selected for downstream analysis. Initial clustering was performed using the Leiden algorithm across all cells. Clusters with high expression of KRT8 were annotated as cancer cells, while those expressing DCN were annotated as CAFs. To integrate all cells, the scPoli method (De Donno et al., 2023) was applied using the initial cell type annotations as input. The following scPoli parameters were used: early_stopping_metric = val_prototype_loss, mode = min, threshold = 0, patience = 20, reduce_lr = enabled (lr_patience = 13, lr_factor = 0.1), n_epochs = 50, pretraining_epochs = 40, eta = 10, and alpha_epoch_-anneal = 100. After integration, clustering was repeated using the Leiden algorithm with a resolution parameter of 0.7. Marker genes for each cluster were identified using the Wilcoxon rank-sum test. Comparison tumoroid scRNA-seq with PDAC atlas To compare the tumoroid clusters with the integrated PDAC atlas, the mean gene expression profile of each tumoroid cluster was computed, as well as the mean expression profiles of cancer and CAF cell types in the PDAC atlas. Pearson correlation coefficients were then calculated between the mean expression profiles of tumoroid clusters and those of the PDAC atlas cell types.

### MERSCOPE experiments

Tumoroids for the spatial transcriptomics are generated in a 96-well plate as described above. After 7 days of co-culture, the tumoroids were fixed with RNAse-free 4% paraformaldehyde (PFA) for 1 hour at room temperature. The PFA was prepared by diluting a 16% PFA stock (Thermo Scientific, 28908) with RNAse-free water (Invitrogen, 10977-035). Fixed tumoroids were washed twice with 1x RNAse-free PBS (Thermo Fisher, AM9625). Fifteen tumoroids were embedded in O.C.T. compound (Tissue-Tek) within a single Cryomold and snap-frozen on dry ice. Samples were stored at −80°C until the MERFISH experiment.

The MERFISH gene probe panel was designed based on the scRNA-seq data from the tumoroid. Genes were selected to represent each cell type and the extracellular matrix. The total number of genes in the panel for this experiment was 300 genes.

RNA detection quality of the samples was qualitatively assessed using the Merscope verification kit. The raw images were opened in the Merscope visualizer software and checked for brightness and abundance of the detected transcripts as well as the background of the tissue. Multiplexed Error-Robust Fluorescence in situ Hybridization (MERFISH) was performed based on the manufacturer’s instruction (“User Guide for Fresh and Fixed Frozen Tissue Sample Preparation, Rev D”). Prior to sectioning, the Merscope slides were treated with 0.1mg/ml poly-D-Lysine (Gibco, A38904-01) for 2h at room temperature and washed with nuclease-free water and let air dry. The OCT block with the embedded tumoroids was then cut with 10-µm thickness using the NX70 Cryostat with a cutting temperature of −16°C/−18°C. Multiple sections were placed on one Merscope slide that was dried for 20min in the cryostat, followed by 30min on a heating plate set to 50°C. After washing off the OCT with nuclease-free PBS the tumoroids were permeabilized in 70% EtOH overnight in a 6 cm Petri dish. Photobleaching was performed with the Merscope photobleacher for 3 hours. Antibody staining was performed using the protocol and antibodies from Vizgen (10400118). The gene probe was incubated for 40h at 37 °C. Gel embedding was performed based on the protocol from Vizgen with Ammonium persulfate (Sigma Aldrich, 09913-100g) and TEMED (Millipore-Sigma, T7024-25ML). Next, the sample was cleared for 24h at 47°C and visually inspected to ensure that the tumoroids were not visible in the gel anymore. Then it was kept at 37°C before starting the run on the instrument. No digestion was performed for the tumoroid experiment. Before the sample was loaded on the Merscope instrument, the section was stained with DAPI and polyT. The spatial transcriptomic data was acquired on the Merscope instrument according to the protocol (“Instrument User Guide”). Four regions of interest were manually drawn around the tumoroids and the data processing of the raw images was performed with the integrated Merscope pipeline on the instrument. To assess the quality of the run the result was inspected with the Vizgen Merscope Visualizer software.

### MERSCOPE data cell segmentation

We performed spatial transcriptomics analysis using MERSCOPE data to characterize gene expression at single-cell resolution within organoid samples. Image preprocessing began with maximum intensity projection (MIP) across Z-stacks for each stain, producing 2D representations of spatial channels. A composite image was generated from selected stains (e.g., PolyT, Cellbound3) for downstream segmentation. Organoids were localized and segmented using a combination of Gaussian filtering, Otsu thresholding, and morphological operations, followed by region labeling and optional region expansion. The top organoid regions were selected by size, and cellular segmentation within these regions was performed using the pretrained Cellpose model (Stringer and Pachitariu, 2025) with adjusted diameter and threshold parameters optimized for our imaging conditions. Transcript coordinates from the MERSCOPE-detected transcript file were converted from global to pixel coordinates using metadata-derived spatial boundaries and a predefined micron-to-pixel conversion factor. Transcripts were assigned to segmented cells by aligning coordinates with labeled masks and counted per cell and gene. Spatially resolved expression matrices and corresponding transcript coordinates were saved for downstream analysis. Additionally, visual overlays, segmentation masks, and cropped image regions for each stain were saved to facilitate quality control and visualization.

### MERSCOPE data analysis

Spatial transcriptomic data generated by MERSCOPE were processed using Scanpy (Wolf et al., 2018). Count matrices and metadata from all distinct tissue regions were loaded, assembled, and merged into AnnData objects. Cells were filtered based on transcript count (>100) and physical area (1,000-20,000 µm^2^), and genes annotated as control probes were removed. To correct for variability in cell size, expression values were normalized by cell volume, applying a scaling factor derived from each cell’s measured area. Following volume adjustment, data were normalized per cell and log-transformed. PCA was performed using the top 100 components. A neighborhood graph was constructed using 15 nearest neighbors and 15 principal components. UMAP was then applied for dimensionality reduction and visualization. Leiden clustering was conducted with a resolution of 0.5 to identify transcriptionally distinct spatial cell populations. Region identity was retained for each cell to enable downstream stratified analyses. To investigate spatial relationships between cell clusters, spatial neighborhood graphs were computed using the spatial_neighbors function from Squidpy (Palla et al., 2022), based on spatial coordinates embedded in the AnnData object. Neighborhood enrichment analysis was then performed using the nhood_enrichment function with Leiden cluster assignments to quantify enrichment or depletion of interactions between clusters, revealing spatial organization patterns within the tissue. Results from the neighborhood enrichment analysis were visualized using the nhood_enrichment plotting function from Squidpy.

To compare transcriptional profiles of spatially resolved cell clusters derived from MERSCOPE data with the reference PDAC atlas, we computed average gene expression per cluster using the Leiden clustering results. Similarly, bulk expression profiles for annotated cell types in the reference PDAC atlas were generated. The comparison was restricted to genes shared between datasets. Pearson correlation analysis was performed between MERSCOPE cluster profiles and reference cell type profiles to assess similarity.

## Supplementary Information

**Extended Data Figure 1.**
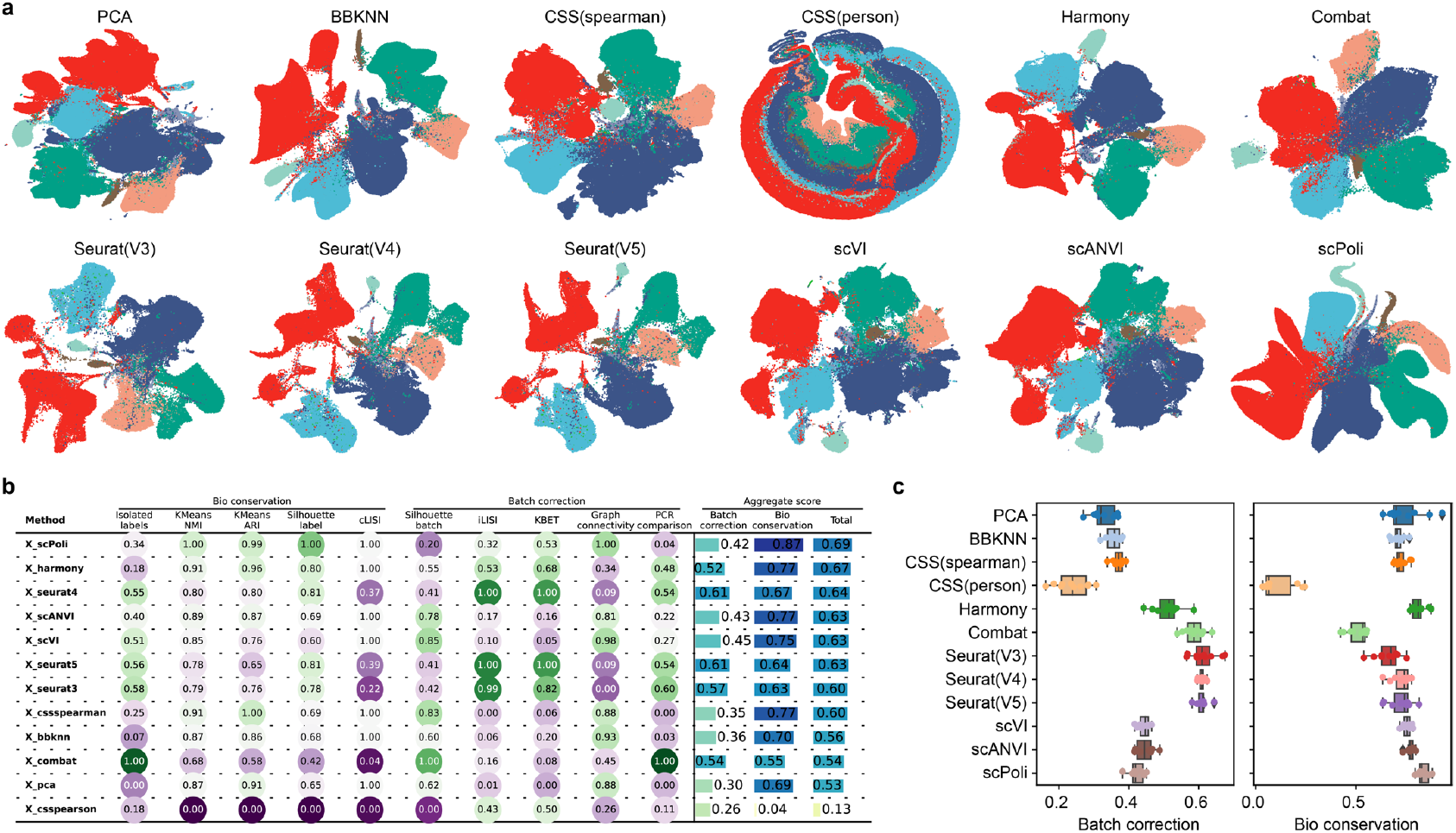
Evaluation of PDAC cell atlas scRNA-seq integration methods. (a) UMAP of tested integration methods and without any data integration (PCA). Dots in all UMAP embeddings are colored by the level 1 cell type annotation. (b) Example scIB benchmarking metrics for all tested integration methods. (c) The boxplot displays the benchmarking results for ten scIB batch correction (left) and biology conservation (right) tests.

**Extended Data Figure 2.**
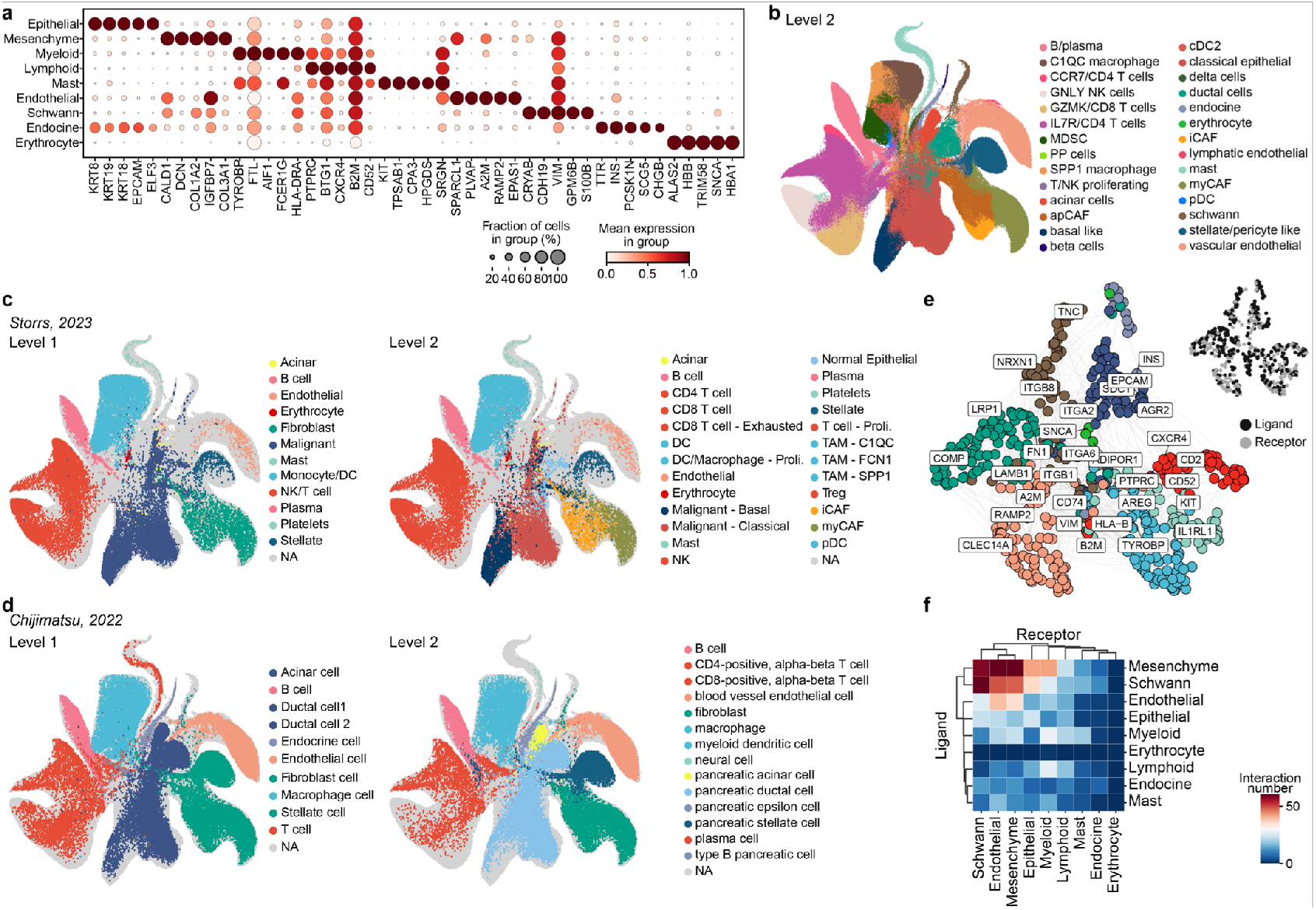
Extended analysis of the integrated PDAC cell atlas. (a) Dotplot showing the top marker genes for each Level 1 cell type as defined in Fig. 1b. (b) UMAP embedding of the integrated atlas annotated with Level 2 cell type identities. (c) UMAP embedding of the integrated atlas with Level 1 (left) and Level 2 (right) annotations as shown in Storrs et al., 2023. (d) UMAP embedding of the integrated atlas with Level 1 (left) and Level 2 (right) annotations as shown in Chijimatsu et al., 2022. (e) Network diagram of inferred ligand-receptor (LR) interactions between cell types in primary tissues, colored by cell type (left) and molecular function (right). (f) Heatmap showing the strength of inferred LR interactions between cell types in primary tissues.

**Extended Data Figure 3.**
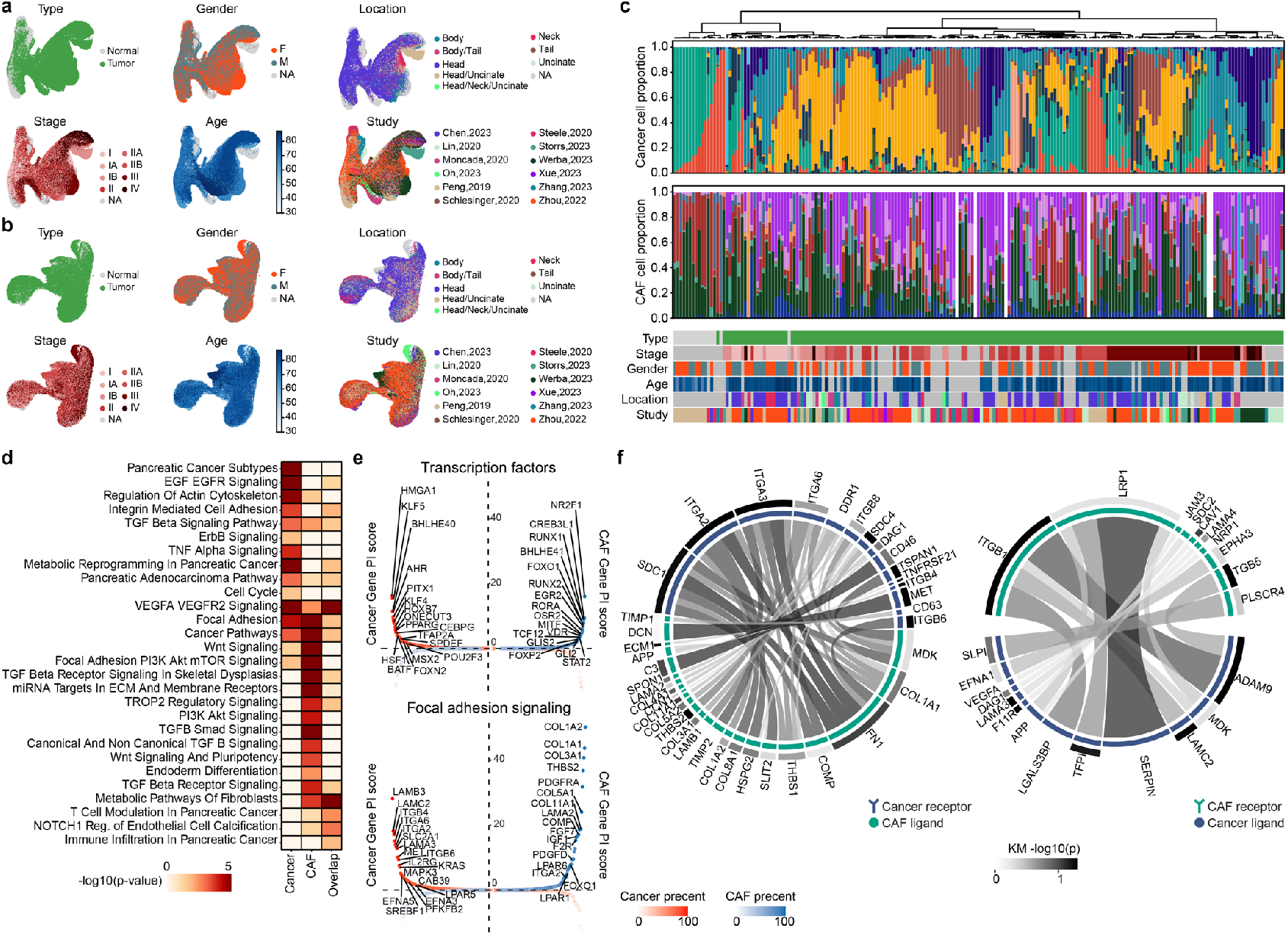
Extended analysis of PDAC cancer and CAF cell sub-atlas. (a) UMAP embedding of cancer cells colored by sample metadata. From left to right and top to bottom, panels represent sample type, gender, anatomical location, tumor stage, age group, and study origin. (b) UMAP embedding of CAF cells colored by the same sample metadata categories as in (a). (c) Stacked bar plots showing the proportions of cancer cell types (top) and CAF subtypes (bottom) across individual samples. Sidebars indicate sample annotations corresponding to the metadata shown in (a) and (b). (d) Heatmap showing key pathways associated with differentially expressed genes specific to cancer cells, CAFs, and genes shared between both populations (as defined in Fig. 2d). (e) Dot plots of all transcription factor genes (left) and CAF-enriched genes involved in focal adhesion and PI3K/Akt/mTOR signaling pathways (right). Genes are sorted left-to-right by cancer gene prioritization (PI) scores and right-to-left by CAF PI scores. Color indicates the percentage of samples with high gene expression. (f) Ribbon plots illustrating ligand-receptor (LR) interactions from CAFs to cancer cells (left) and from cancer cells to CAFs (right) in the PDAC primary atlas. Ribbon color and width represent interaction strength across all cancer and CAF subtypes. Outer scale bars indicate the−*log*_10_(*p*) values from survival analysis.

**Extended Data Figure 4.**
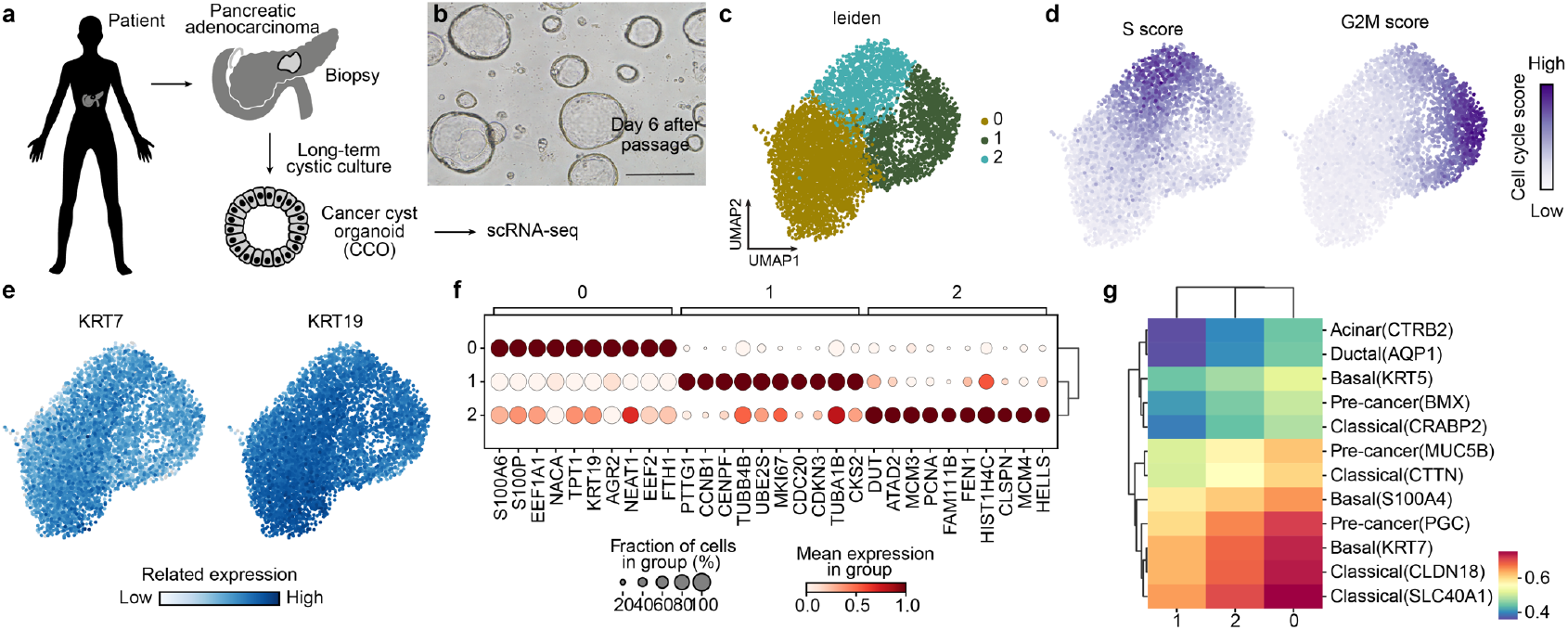
Single-cell transcriptome analysis of pancreatic cancer cyst organoid cultures. (a) Cancer cyst organoid (CCO) lines were established from biopsies of PDAC patients. CCOs were expanded over multiple passages, and scRNA-seq was performed on cells collected at day 7 post-passage. Scale bar: 400 µm. (b) Brightfield image of CCOs in 3D Matrigel culture 6 days after passage. (c) UMAP embedding of CCO scRNA-seq data, colored by cell cluster. (d) UMAP embedding of the same dataset, colored by cell cycle state: S phase (left) and G2/M phase (right). (e) Feature plots showing the expression of PDAC cancer cell markers KRT19 and KRT17. (f) Dotplot displaying the top 10 marker genes for each cluster identified in (c). (g) Heatmap showing the correlation between scRNA-seq-defined CCO clusters and primary cancer cell types from the atlas in Fig. 2b.

**Extended Data Figure 5.**
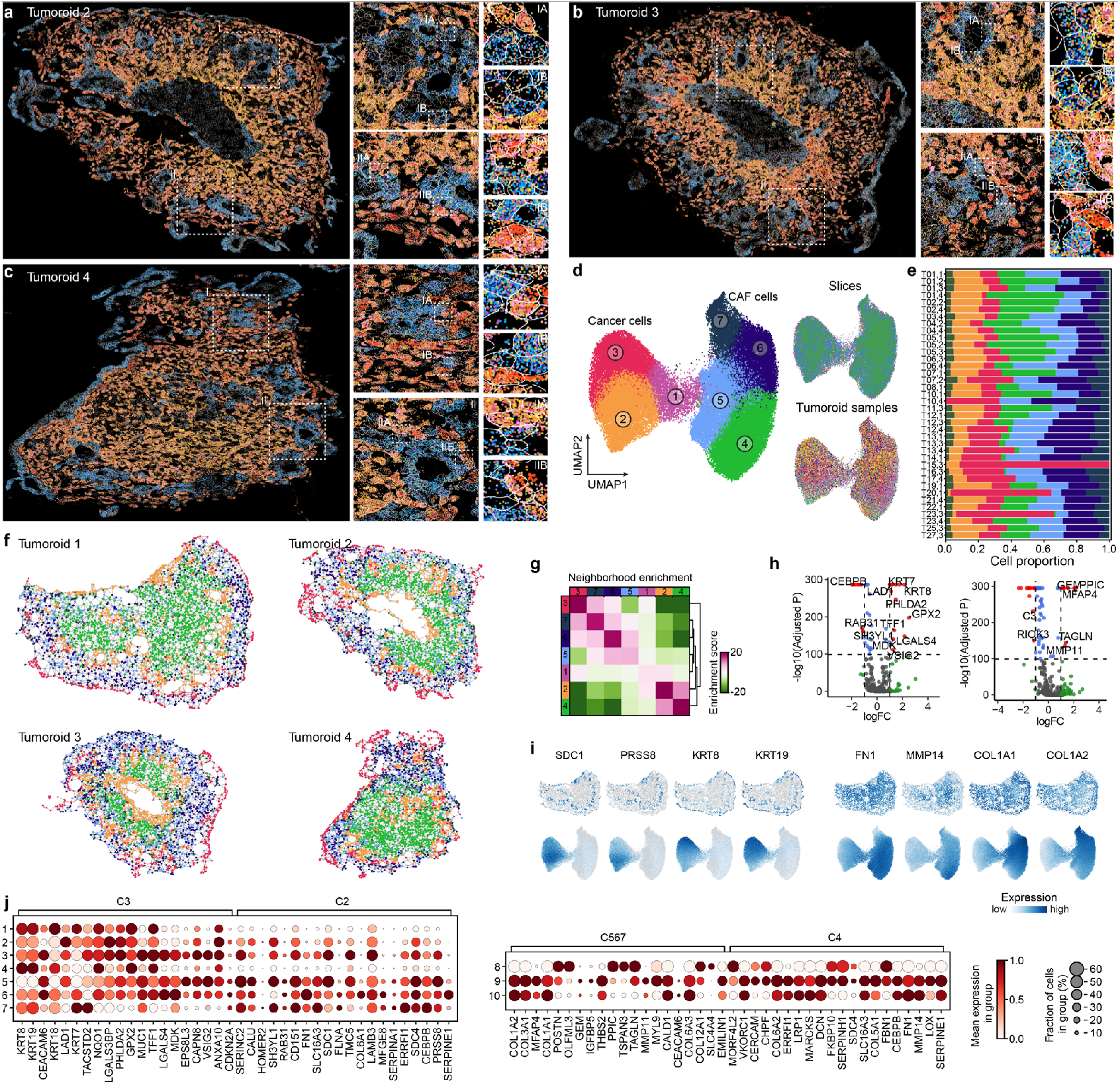
Extended analysis of tumoroid spatial transcriptome data. (a-c) Spatial transcriptomic maps of three representative tumoroids showing four gene transcript clusters corresponding to those in Fig. 3e. (d) UMAP embedding of all PDAC tumoroid spatial transcriptomics samples, colored by Leiden cluster (top), sample slice (bottom left), and tumoroid identity (bottom right). (e) Cell-type proportions across all tumoroid samples, with cluster colors matching those in (d). (f) Spatial distribution of clusters in four representative tumoroids, colored as in (d). (g) Heatmap showing neighborhood enrichment across all spatial clusters. (h) Volcano plots showing differentially expressed genes between clusters 2 vs. 3 (left) and clusters 4 vs. 5/6/7 (right), with selected marker genes highlighted. (i) Spatial (top) and UMAP (bottom) views showing the expression of representative genes identified in (h). (j) Dot plots showing the top 20 differentially expressed genes: left, for clusters 2 vs. 3 mapped to scRNA-seq cancer clusters from Fig. 3c; right, for clusters 4 vs. 5/6/7 mapped to scRNA-seq CAF clusters from Fig. 3c.

**Extended Data Figure 6.**
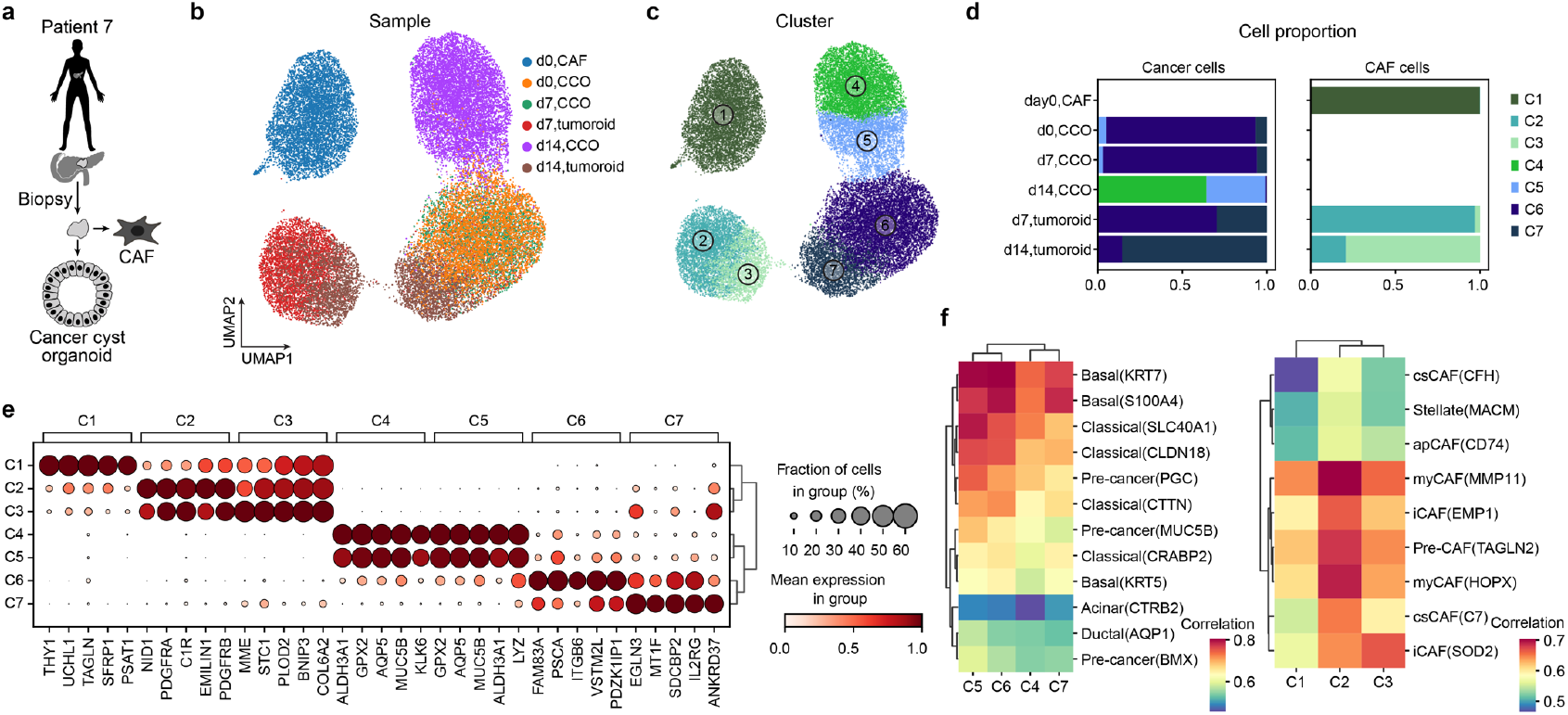
Single-cell transcriptome analysis of autologous cancer-CAF tumoroids. (a) Schematic representation of sample collection for autologous PDAC in vitro models. (b-c) Integrated UMAP embedding highlighting identified samples (b) and clusters (c). (d) Bar plot summarizing the proportions of each cell cluster across all samples. (e) Dotplot showing normalized expression levels of top marker genes for each identified cluster. (f) Heatmap displaying correlation between cancer cell clusters (left) and CAF clusters (right) identified by scRNA-seq in tumoroids and primary cell types derived from the atlas in Fig. 2b,c.

**Extended Data Figure 7.**
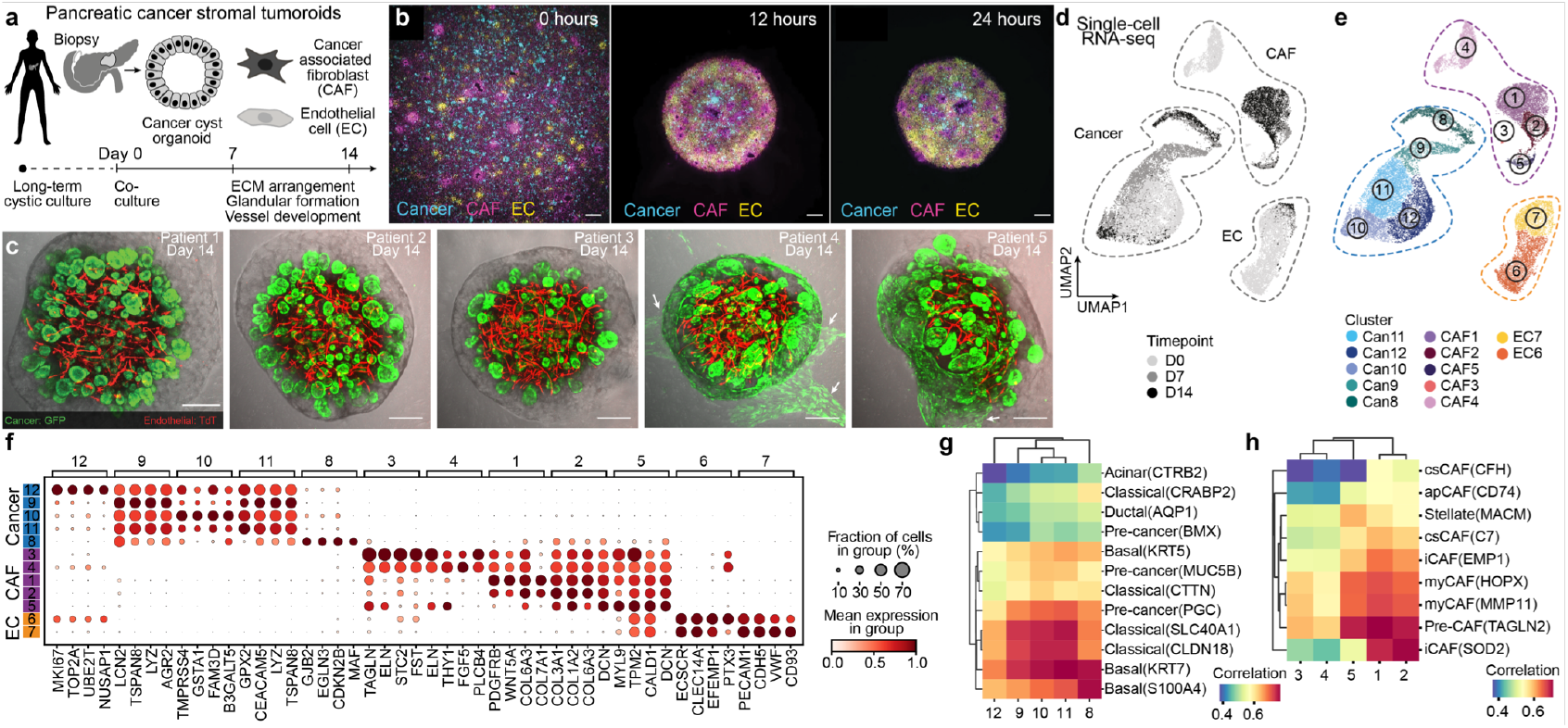
Assembly of stroma-rich tumoroids. (a) Long-term cancer cyst organoid (CCO) cultures derived from patients with pancreatic adenocarcinoma (PDAC) co-cultured in a 3D matrix with endothelial cells (ECs) and cancer-associated fibroblasts (CAFs), self-organizing into a complex tumoroid microenvironment. Over 14 days, fibrous connective tissue forms, vessels sprout and organize, and cancer cells develop 3D glandular structures within multilineage tumoroids. (b) Imaging of reporter-labeled cancer cells (teal blue), CAFs (pink), and ECs (yellow) at 0, 12, and 24 hours after co-culture. Scale bar: 10 µm. Representative image of a Day-14 tumoroid using cancer cells from five different patients and ECs stably expressing EGFP and TdTomato. arrow: EGFP+ migrated cancer cells. Scale bar: 250 µm. (d-e) scRNA-seq performed on input cells in mono-culture (Day 0) and tumoroids after 7 and 14 days of co-culture. UMAP cell embedding of scRNA-seq data is shown, colored by time point (d) and by cluster (e). (f) Heatmap showing the normalized expression of cluster marker genes. (g) Heatmap depicting the correlation between cancer cell clusters (left) and CAF clusters (right) identified by scRNA-seq in tumoroids and primary cell types from the atlas in Fig. 2b,c.

**Extended Data Figure 8.**
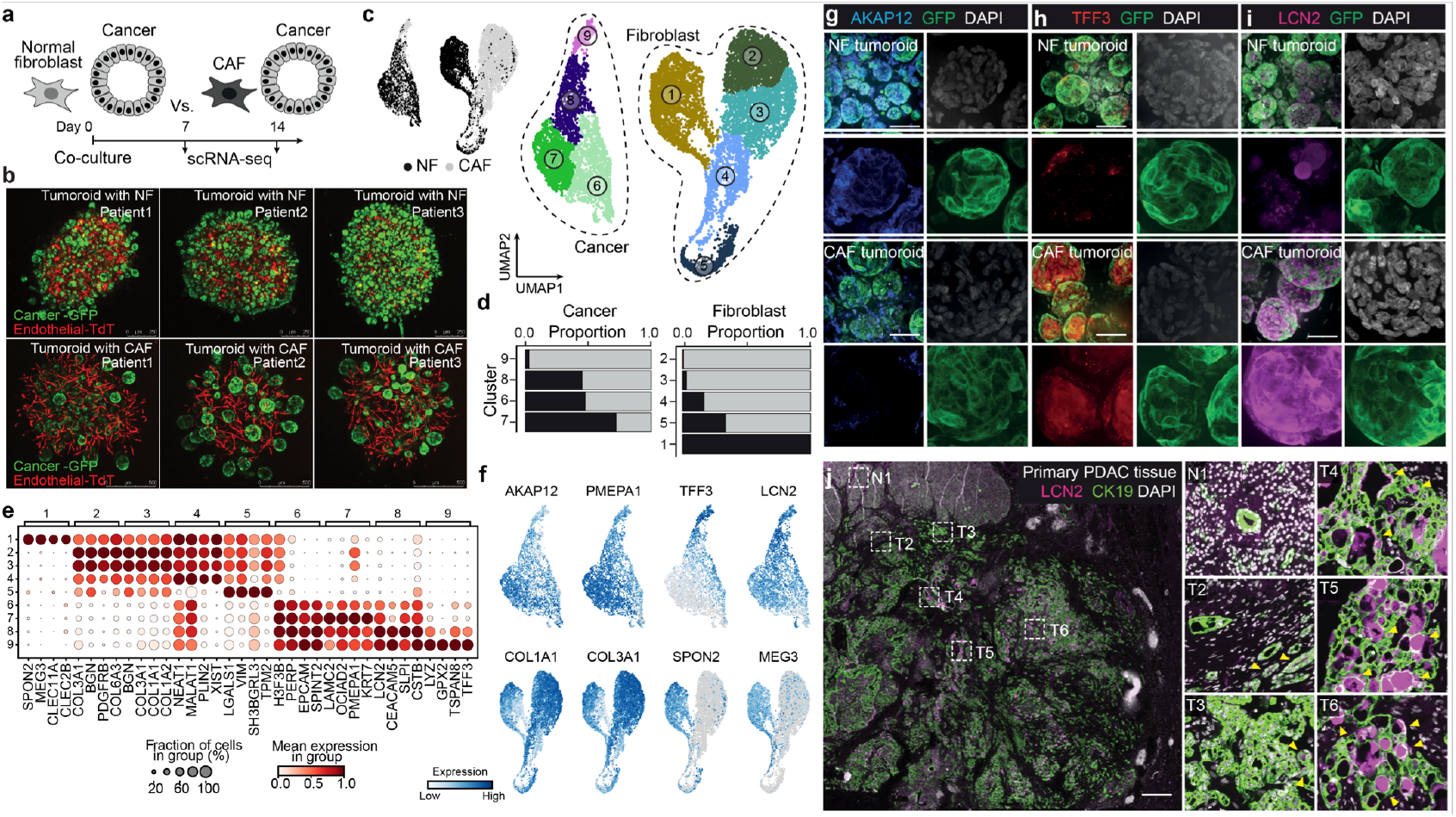
Comparison of tumoroids with normal or cancer associated fibroblasts. (a) Tumoroids containing normal fibroblasts (NFs) or cancer-associated fibroblasts (CAFs) were generated and analyzed by scRNA-seq. (b) Representative images of tumoroids generated with NFs (A, upper row) or CAFs (B, lower row), with endothelial cells labeled with TdTomato and cancer cells labeled with EGFP. Scale bar: 250 µm. (c) UMAP embedding of scRNA-seq data, colored by tumoroid type (left) and cluster (right), with cancer and fibroblast cells encircled and labeled. (d) Stacked bar plot illustrating the proportion of cancer and fibroblast cells per cluster, colored by tumoroid type. (e) Dotplot showing normalized expression levels of cluster marker genes. (f) UMAP embedding colored by expression of feature genes. (g-i) Immunofluorescence staining for AKAP12 (g), TFF3 (h), and LCN2 (i) protein expression in NF and CAF tumoroids. Cancer cells stably express GFP, and nuclei are labeled with DAPI (white). Scale bar: 100 µm. (j) Immunofluorescence of LCN2 and CK19 in Primary PDAC tissue. Insets show 1 location within non-cancerous pancreas tissue (N1) as well as 5 locations within the tumor (T2-6). Nuclei stained with DAPI (white). Scale bar: 1mm.

**Extended Data Figure 9.**
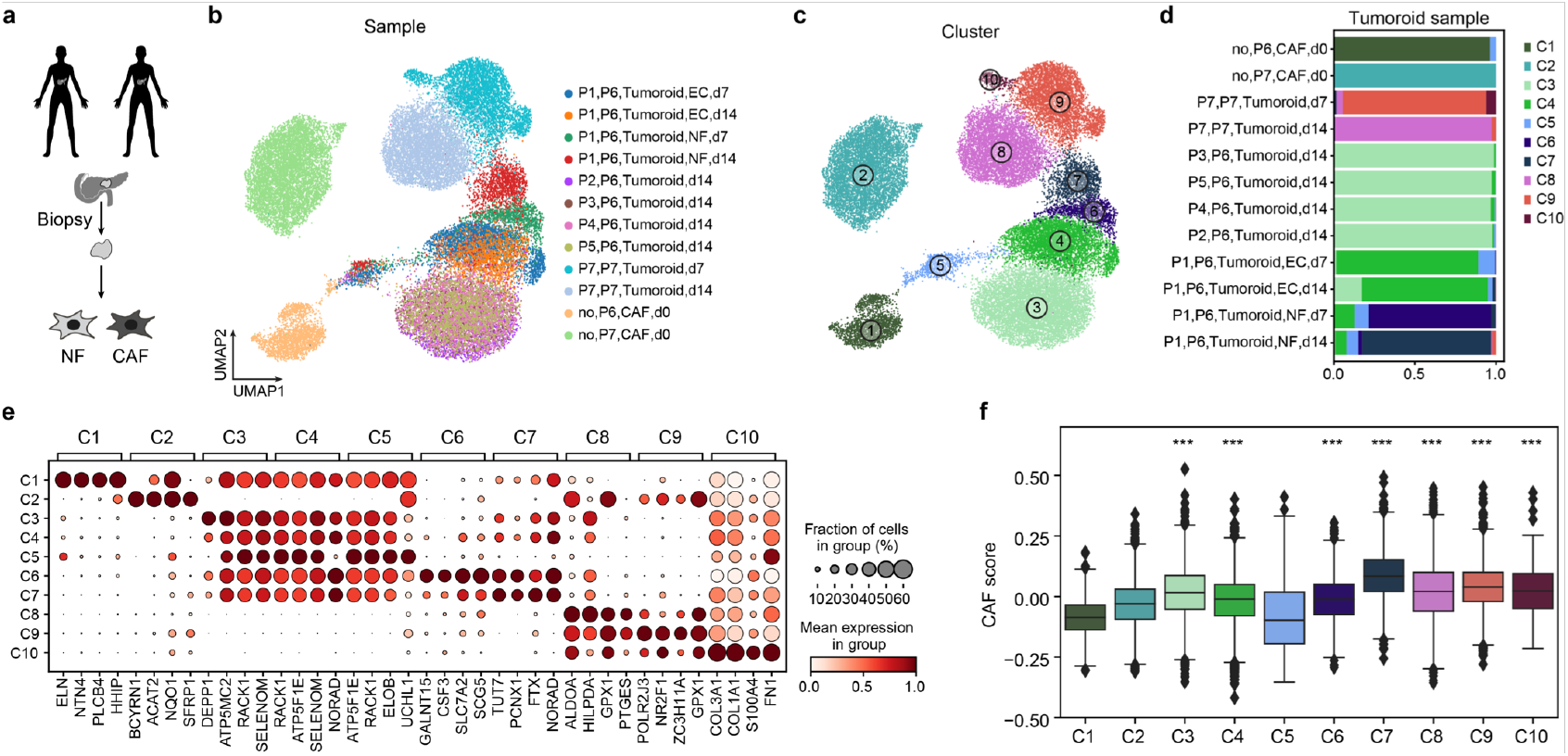
Integrated analysis of tumoroid fibroblasts. (a) Schematic representation of sample collection for autologous PDAC in vitro models across different patients and fibroblast types (normal fibroblasts [NFs] or cancer-associated fibroblasts [CAFs]). (b-c) Integrated UMAP embedding of all fibroblasts, highlighting individual samples (b) and identified clusters (c). (d) Barplot summarizing the proportions of each cell cluster across all samples. (e) Dotplot showing normalized expression levels of top marker genes for each identified cluster. (f) Boxplot showing CAF gene scores across all clusters with Mann-Whitney U test p-values comparing each cluster (C3-C10) to merged C1 and C2 (^*^p < 0.05, ^**^p < 0.01, ^***^p < 0.001).

**Extended Data Figure 10.**
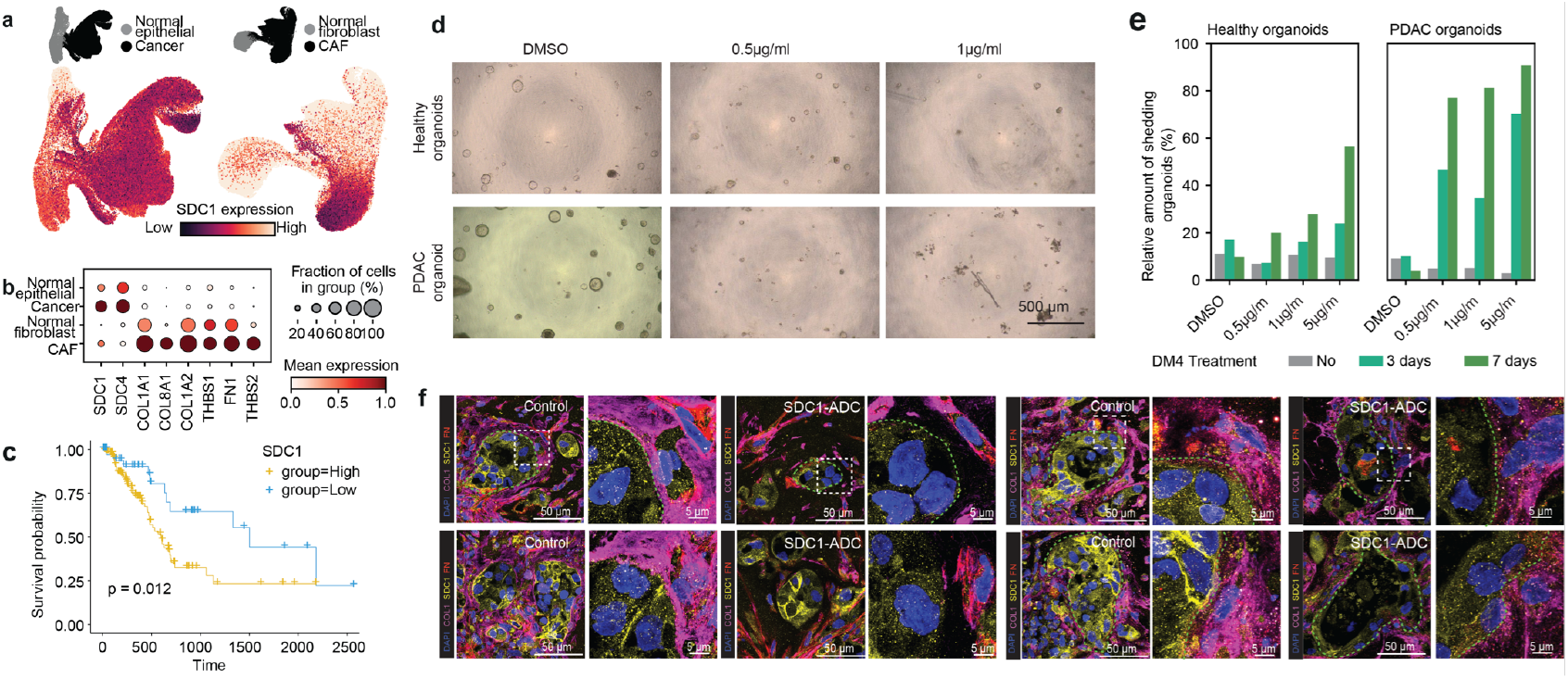
Additional analyses demonstrating SDC1 is a therapeutic target regulating tumoroid growth. (a) UMAP embedding of the integrated PDAC epithelial (left) and mesenchymal (right) atlas from Fig. 2b,c, highlighting cancer and cancer-associated fibroblast (CAF) cells. The bottom UMAP displays SDC1 expression in the PDAC epithelial (left) and mesenchymal (right) atlas. (b) Dot plot illustrating the expression of selected ligand and receptor genes (as shown in Fig. 4a) in the PDAC cell atlas from Fig. 2b,c. (c) Kaplan-Meier survival analysis of PDAC patients stratified by high and low SDC1 expression. (d) Representative brightfield images of healthy and PDAC-derived organoids treated with DMSO or DM4 (0.5 µg/ml and 1 µg/ml) for 7 days. (e) Quantification of organoid formation efficiency in healthy and PDAC-derived organoids treated with DMSO or DM4 for 7 days. (f) Representative immunofluorescence images of PDAC tumoroids stained for DAPI (blue), SDC1 (yellow), COL1 (magenta), and FN (red) under control and DM4 treatment conditions. Scale bar, 50 µm. (g) Representative immunofluorescence images of PDAC tumoroids stained for DAPI (blue), SDC1 (yellow), TNC (magenta), and THBS1 (red) under control and DM4 treatment conditions. Scale bar, 50 µm.

